# Living with high potassium: a balance between nutrient acquisition and stress signaling during K-induced salt stress

**DOI:** 10.1101/2021.07.01.450778

**Authors:** Pramod Pantha, Dong-Ha Oh, David Longstreth, Maheshi Dassanayake

**Affiliations:** Department of Biological Sciences, Louisiana State University, Baton Rouge, LA 70803, USA

**Keywords:** abiotic stress, antagonistic pleiotropy, molecular phenotype, multi-omics, nutrient transport, osmoprotectants and antioxidants, potassium-induced salt stress, systems biology

## Abstract

- High potassium (K) in the growth medium is more toxic to plants than Na at similar concentrations. However, the molecular mechanisms underlying plant responses to K-induced salt stress are virtually unknown.
- We examined *Arabidopsis thaliana* and its extremophyte relative *Schrenkiella parvula*, using a comparative multi-omics approach to identify cellular processes affected by excess K and understand which deterministic regulatory pathways are active to avoid tissue damage while sustaining growth.
- *A. thaliana* showed limited capacity to curb excess K accumulation and prevent nutrient depletion contrasting to *S. parvula* which could limit excess K accumulation without restricting nutrient uptake. Facilitated by a targeted transcriptomic response, promoting nitrogen uptake along with other key nutrients and uninterrupted N assimilation into primary metabolites during excess K-stress allowed *S. parvula* to boost its antioxidant and osmolyte pools concurrently leading to sustained growth. Antithetically, *A. thaliana* showed transcriptional responses indicative of a poor balance between stress signaling, increased ROS levels, and reduced photosynthesis, subsequently leading to inhibited growth.
- The ability to regulate independent nutrient uptake and a coordinated transcriptomic response to avoid non-specific stress signaling are two main deterministic steps towards building stress resilience to excess K^+^-induced salt stress.

## Introduction

Can excess potassium (K^+^) in soil be too much of a good thing for plants? Its role as an essential macronutrient for plants is well established (Wang & Wu, 2013). The cytosolic concentration of K^+^ around 100 mM is typically higher than soil concentrations found in most agricultural soils (Maathuis, 2009). Past studies have focused K^+^ uptake into plants from low concentrations in the soil. Consequently, we barely understand excess K^+^-induced salt stress in plants. While toxicity in plants exposed to high concentrations of nutrients such as boron (Wang *et al.*, 2021), copper (Lequeux *et al.*, 2010), and nitrogen (Yoshitake *et al.*, 2021), have been investigated, the molecular mechanisms behind K^+^ toxicity are virtually unknown.

Worldwide there are many soils naturally high in K^+^ (Duval *et al.*, 2005; Warren, 2016). Many industrial as well as agricultural processing plants produce wastewater with exceptionally high K^+^ concentrations considered excessive for plant growth (Arienzo *et al.*, 2009). During wastewater treatments, unlike N, P, or organic matter which are typically processed using microbial activity, K is concentrated due to evaporation. The need to use alternative agricultural lands and recycled wastewater for irrigation is a current necessity (IPCC, 2021). These needs cannot be addressed without foundational knowledge on how plant nutrient balance is achieved in the absence of prime agricultural land. Therefore, knowing the tolerance mechanisms against high K becomes an impending need in our quest to convert marginal lands into productive agricultural lands.

Past studies have reported that excess K^+^ severely affect growth of multiple crops and even halophytes (Eijk, 1939; Ashby & Beadle, 1957; Eshel, 1985; Matoh *et al.*, 1986; Wang *et al.*, 2001; Ramos *et al.*, 2004; Richter *et al.*, 2019; Zhao *et al.*, 2020). These studies suggest that K^+^ may not elicit the same physiological or metabolic stresses Na^+^ does and plants may require distinct genetic pathways in addition to canonical salt response pathways (Pantha & Dassanayake, 2020) to survive high K-induced salt stress.

In this study, we aimed to identify K-induced salt stress responses and deduce the underlying cellular mechanisms plants have evolved to adapt to K-toxicity. We compared *Arabidopsis thaliana*, sensitive to high K^+^, to its extremophyte relative, *Schrenkiella parvula* that thrives in high K^+^ soils (Nilhan *et al.*, 2008; Oh *et al.*, 2014). We examined stress responses exhibited by the two model species to high K^+^ treatments using a multi-omics approach to identify the relevant genetic pathways influential in regulating or are affected by K-toxicity. Our results revealed an extensive ionomic, metabolic, and transcriptomic reprogramming during high potassium stress in the stress-sensitive model while providing novel insights into how the extremophyte model had evolved alternative metabolic and transcriptomic adjustments to enable growth under excess potassium.

## Materials and methods

Detailed methods in Supporting Information Methods S1.

### Plant Material

*Arabidopsis thaliana* (Col-0) and *Schrenkiella parvula* (Lake Tuz) were grown in controlled environments hydroponically, on plates, or in soil at 23°C, 12/12 h light/dark, and 60% relative humidity (Wang *et al.*, 2021). Multi-omics data were generated from plants grown in 1/5th Hoagland’s solution which included 1.2 mM K^+^ to support plant growth in all conditions. Supplemental KCl (150 mM) treatments provided excess K^+^ beyond its expected range serving as a nutrient (0.1-6 mM) (Ashley *et al.*, 2006). Tissues were harvested at 0, 3, 24, and 72 hours after treatment (HAT) for ionomic, metabolomic, and transcriptomic profiling (Fig. S1).

### Physiological assessments

Root growth assessments were made using plate-grown seedlings scanned and processed with ImageJ (Ferreira & Rasband, 2012). Reproductive stage assessments were taken from soil grown plants treated with 0-400 mM NaCl or KCl given for at least two weeks. CO_2_ assimilation rates were measured on hydroponically grown plants using an infrared gas analyzer (LI-6400XT, LI-COR, Lincoln, USA).

### Elemental profiling

Samples were processed following Ziegler et al. (2013) to quantify K, P, Mg, S, Ca, Al, Zn, Co, Ni, Fe, Se, Cu, B, Mn, Mo, As, Rb, Cd, Na, Li, and Sr via inductively coupled plasma mass spectrometry (Baxter *et al.*, 2014). Carbon and nitrogen were quantified using a Costech 4010 elemental analyzer (Costech, Valencia, USA). Significant differences were calculated based on one-way ANOVA, followed by Tukey’s post hoc test (p≤0.05) using agricolae and visualized as heatmaps using pheatmap (R packages, v. 1.0.12) (Kolde, 2012).

### Metabolite profiling

Non-targeted high throughput metabolite profiling was conducted on frozen samples using gas chromatography-mass spectrometry (Fiehn *et al.*, 2008). Significance tests were performed as described for ionomics and visualized using circlize, R (Gu *et al.*, 2014).

### RNA-seq analyses

Total RNA was extracted from triplicates of root and shoot tissues harvested at 0, 3, 24, and 72 HAT with 150 mM KCl (Fig S1). Strand-specific RNA-seq libraries were sequenced on Illumina HiSeq4000. Reads uniquely mapped to *A. thaliana* TAIR10 or *S. parvula* (Dassanayake *et al.*, 2011) v2.2 gene models (https://phytozome-next.jgi.doe.gov/) (Table S2) were counted for identifying differentially expressed genes based on DESeq2 (Love *et al.*, 2014) at p-adj ≤0.01. Orthologous gene pairs were identified using CLfinder-OrthNet (Oh & Dassanayake, 2019) and further refined as described in Wang et al. (2021). Median normalized expression of orthologs were used to determine temporal co-expression clusters based on fuzzy k-means clustering (Gasch & Eisen, 2002). Enriched functional groups were identified using BinGO (Maere *et al.*, 2005) (p-adj ≤0.05) and further summarized to non-redundant functional groups using GOMCL (Wang *et al.*, 2020).

### Histochemical analysis

Leaves were stained for H_2_O_2_ and O_2_^-^ using 3,3′- diaminobenzidine and nitroblue tetrazolium (Jabs *et al.*, 1996; Daudi & O’Brien, 2012).

## Results

### KCl is more toxic than NaCl at the same osmotic strengths

Excess K^+^ exerted more severe growth disturbances than observed for Na^+^ at the same concentrations in both species (Figs 1, S2, S3). The extremophyte, *S. parvula*, was more resilient to higher concentrations of KCl than *A. thaliana* before it showed significant alterations to root, leaf, or silique development (Figs 1b-d, S2, S3). CO_2_ assimilation was reduced in *A. thaliana* in response to 150 mM KCl within 24 HAT, corresponding to a reduction in total shoot carbon (Fig. 1e-f). Under long-term treatments, there was a greater reduction in total leaf area in both species with high K^+^ than Na^+^ compared to control conditions (Figs 1d, S3b). These results suggest that despite similar levels of osmotic stress elicited by K^+^ and Na^+^ at equal concentrations, K^+^ may exert additional stresses different from Na^+^ that may initiate unique responses.

**Fig. 1.**
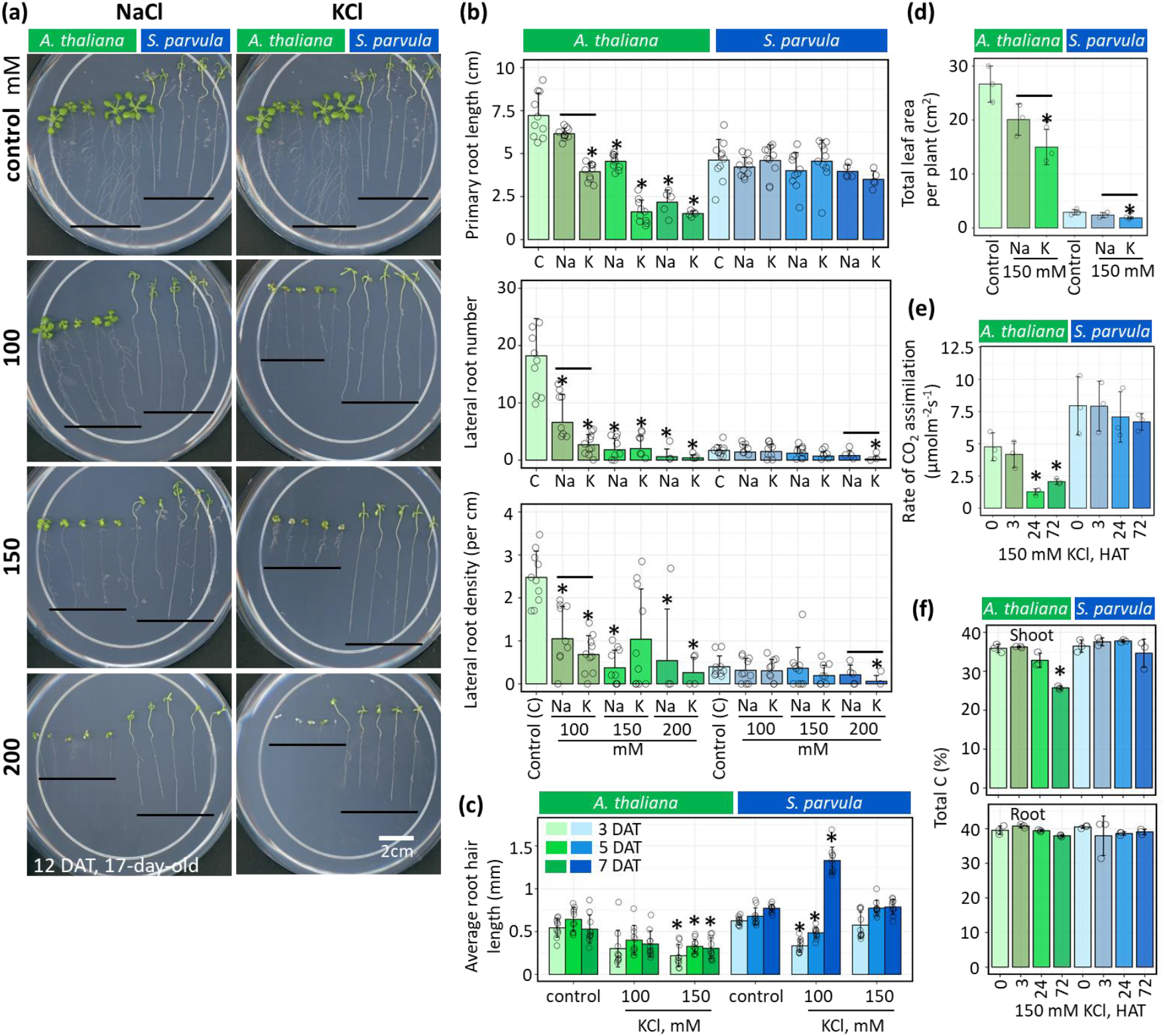
KCl is more toxic than NaCl at the same osmotic strength. (a) Seedlings of *Arabidopsis thaliana* (left on each plate) and *Schrenkiella parvula* (right on each plate) on 1/4^th^ MS media supplemented with 0 to 200 mM NaCl or KCl. Black line on plates mark end of root tips. (b) Primary root length, lateral root number, and lateral root density measured on 17-day-old seedlings grown under conditions used in (a) based on a 12-day treatment of NaCl or KCl. (c) Average root hair length of 10 longest root hairs measured under the same growth conditions used in (a) monitored for a week. (d) Total leaf area per plant measured when plants developed the first floral bud following 1-2 weeks of salt treatments in a hydroponic medium. (e) Photosynthesis measured as the rate of CO_2_ assimilation of the entire shoot/rosette in 25-day-old hydroponically grown plants and monitored up to 72 HAT. (f) Total carbon as a % weight based on total dry mass for root and shoot tissue under conditions used in (e). Minimum 5 plants per condition used in b and c and a minimum of 3 plants per condition used for d and e. Asterisks indicate significant changes between the treated samples to its respective control samples (t-test with p ≤ 0.05). Black lines above bars in (b) and (d) mark the conditions where K causes a severe reduction than Na compared to the control at similar K and Na concentrations. Data are presented as mean of at least 3 independent biological replicates ± SD. Open circles indicate the measurements from each plant (b-c) and replicate (d-f). DAT-Days after treatment, HAT-Hours after treatment.

Overall, 150 mM KCl was sufficient to induce physiological stress responses in *A. thaliana* within a 3-day period, while impacting long-term growth in *S. parvula*. Notably, 150 mM KCl treatments exceeding two weeks were lethal to *A. thaliana* in tested conditions. Therefore, we selected 150 mM KCl for our cross-species comparative-omics study to investigate K^+^ toxicity responses, when both species are expected to show active cellular responses during the early stages of the treatment at 0, 3, 24, and 72 HAT (Fig. S1).

### Excess K^+^ causes nutrient depletion in *A. thaliana* but not in *S. parvula*

Plants have evolved multiple transporters to facilitate K^+^ entry into roots rather than its exit pathways (Shabala & Cuin, 2008). Therefore, we hypothesized that high K^+^ in soils will lead to high K^+^ contents in plants and shoots will accumulate more of the excess K^+^ in the transport sequence of soil-root-shoot. We predicted that the differential tissue compartmentalization of K^+^ will alter K-dependent cellular processes. With elemental profiling of 18 nutrients and six common toxic elements (Methods S1, Fig. S1), we aimed to test if high K^+^ would, (a) accumulate as an isolated process; (b) cause nutrient imbalances; or (c) allow entry of other toxic elements and thereby indirectly cause toxicity symptoms.

Shoot K^+^ levels were higher than in roots, but this difference was strikingly larger in *S. parvula* than in *A. thaliana* under control conditions (Fig. 2a). When treated with excess K^+^, *A. thaliana* initially reduced its root K^+^ levels, but was unable to restrict accumulation with prolonged stress, leading to significant increases by 72 HAT. Importantly, *S. parvula* maintained shoot K^+^ levels comparable to its control during K^+^ treatments (Fig. 2a).

**Fig. 2.**
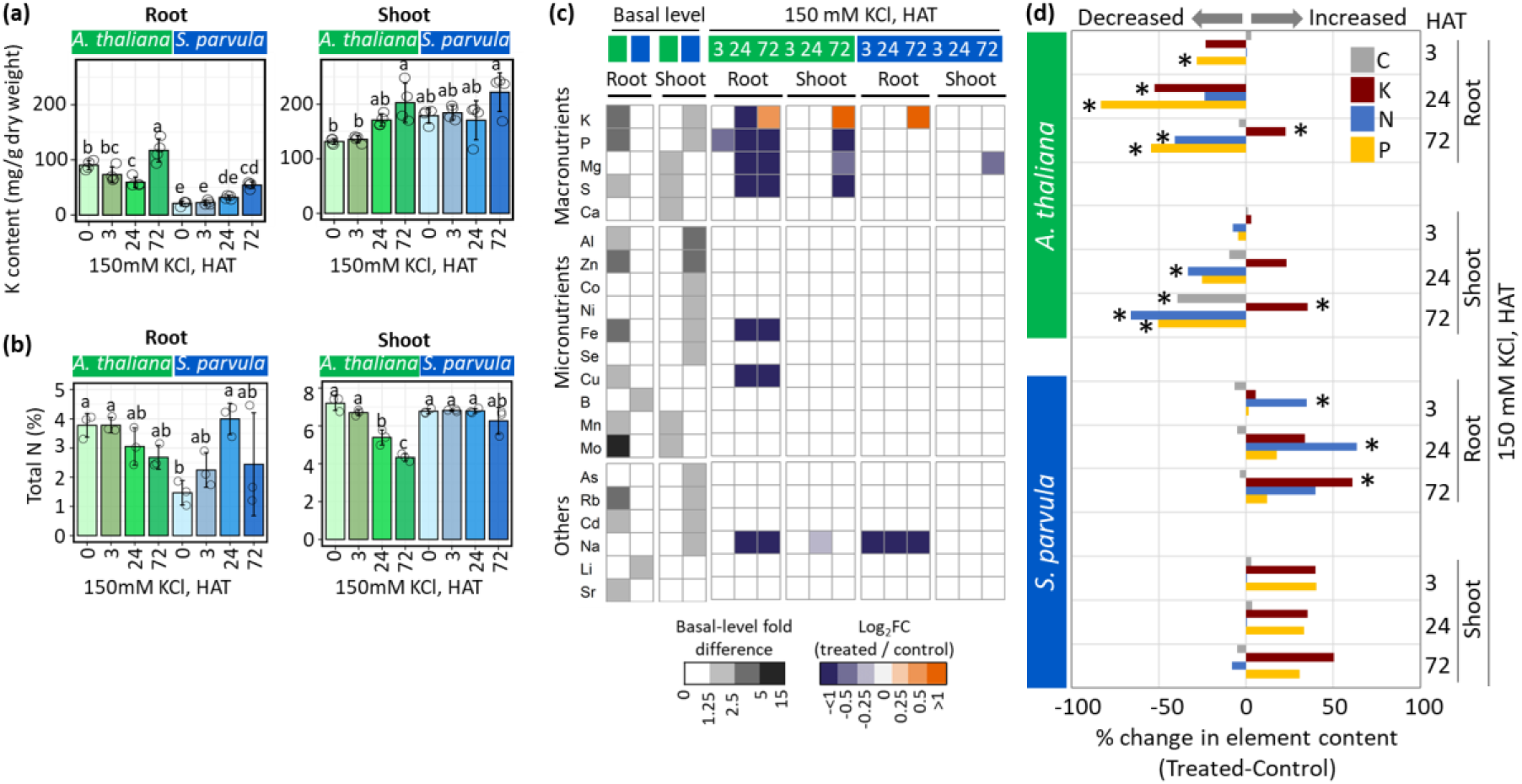
Excess K accumulation cause severe nutrient imbalance in *A. thaliana* compared to *S. parvula.* (a) Total potassium (K) accumulation between *A. thaliana* and *S. parvula*. (b) Total nitrogen (N) levels in *A. thaliana* and *S. parvula*. (c) Ionomic profiles of 15 nutrients and 6 other elements quantified during excess K^+^ treatments. (d) Percent change in CNPK elemental content in roots and shoots of *A. thaliana* and *S. parvula* during excess K^+^ treatments. Data are represented as mean of at least 4 (for a and c) and 3 (for b) independent replicates with ± SD based on 5-8 hydroponically grown plants per replicate. Total elemental compositions reported after normalization for dry weight. Significant differences are based on one-way ANOVA followed by Tukey’s post-hoc test*, p-value* ≤0.05. Asterisks or different letters assigned to bars in a, b, and d indicate significant differences between treatments and control. Open circles indicate data from each replicate. HAT-Hours after treatment.

Potassium accumulation in *A. thaliana* resulted in a severe nutrient imbalance leading to the depletion of seven nutrients including nitrogen (Fig. 2b-d). In line with uninterrupted root growth observed under 150 mM KCl (Fig. 1a-c), *S. parvula* showed a remarkable capacity in maintaining its macronutrients levels (Fig. 2d). Moreover, other elements did not accumulate under excess K^+^ to suggest indirect toxicity effects in either species (Fig. 2c). The overall ionomic profiles suggest that constraining K^+^ accumulation while upholding other nutrient uptake processes is necessary for K^+^ toxicity tolerance.

### *S. parvula* is more responsive than *A. thaliana* at the metabolome level to excess K^+^ stress

We obtained 472 (145 known and 327 unannotated) metabolite profiles across tissues and time points to identify metabolic processes influenced by K^+^ stress (Table S1). The relative abundances of metabolites were highly correlated between the two species for similar conditions (Fig. 3a). For downstream analysis, we only considered the known metabolites with significant abundance changes (MACs) between control and KCl treated conditions (Fig. 3b).

**Fig. 3.**
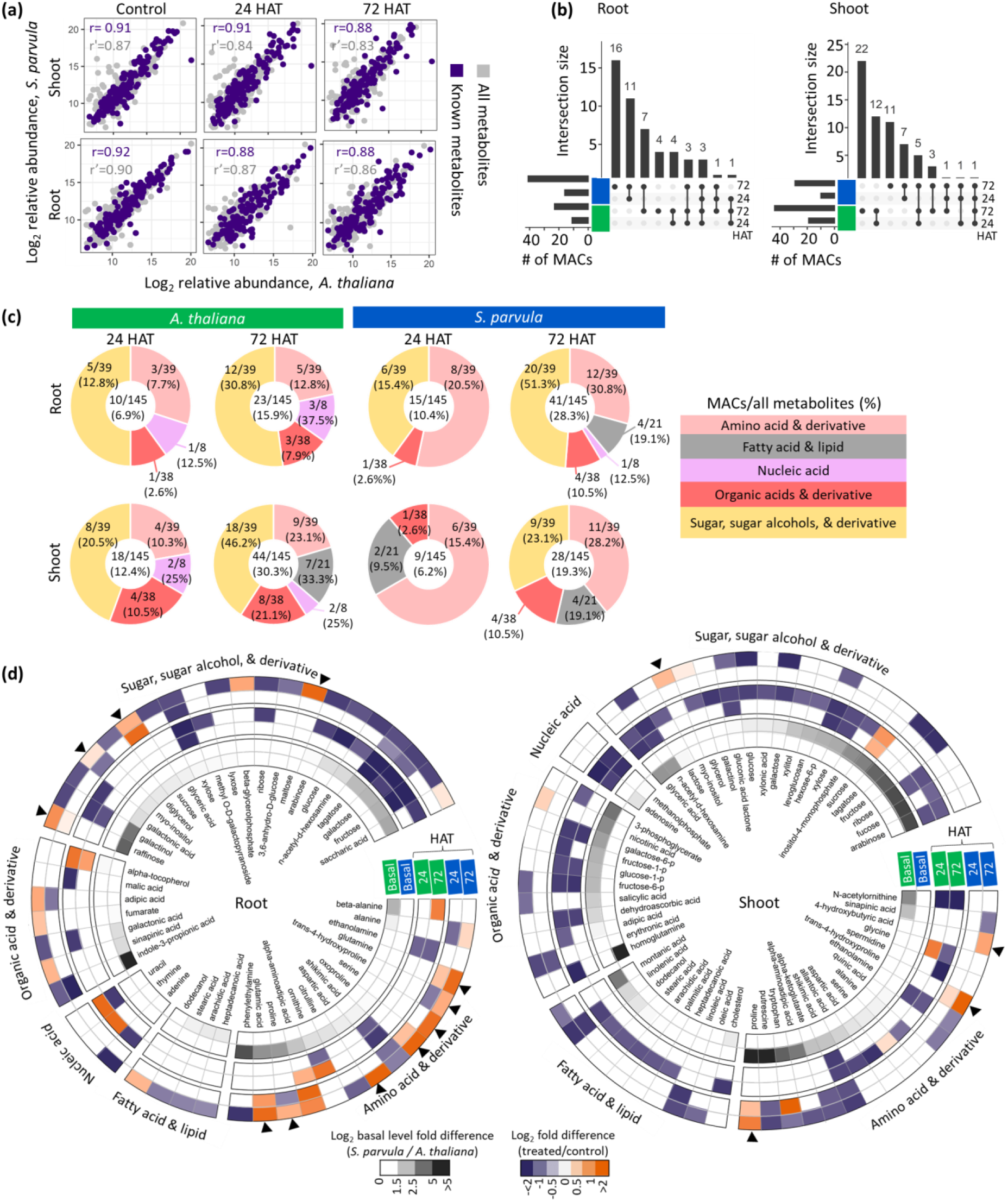
*S. parvula* metabolome is more responsive than *A. thaliana* and induces specific antioxidants and osmoprotectants during excess K^+^ stress. (a) Correlation between *A. thaliana* and *S. parvula* quantified overall metabolome profiles. Pearson correlation coefficient is calculated for 145 known metabolites (r) and all metabolites including the unannotated metabolites (r’). (b) Known metabolites that significantly changed in abundance (MACs) at 24 and 72 hours after treatment (HAT). (c) Overview of the temporal changes in primary metabolite pools based on broad functional groups. The numbers indicate MACs followed by (“/”) the total number metabolites quantified in each group. (d) Metabolites in each functional group mapped to represent their abundance starting at basal level (inner circles) to 24 and 72 HAT. Antioxidants and osmolytes highlighted in Results are marked with arrowheads in the outer circles. Significance tests for metabolite abundance (to detects MACs) were performed with one-way ANOVA followed by Tukey’s post-hoc test*, p-value* ≤0.05. Data presented as mean of at least 3 independent replicates ± SD given using ≥ 5 hydroponically grown plants per replicate. HAT-Hours after treatment.

We observed three distinct trends from metabolite profile comparisons (Fig. 3b-d). First, with longer stress durations, the number of MACs increased in both species. Second, *S. parvula* contained more MACs in roots, even though it had fewer physiological and ionomic adjustments compared to *A. thaliana* under excess K^+^ (Figs 1-2). *A. thaliana* shoots had more MACs than in *S. parvula* consistent with a reduction in photosynthesis seen in *A. thaliana* during K^+^ toxicity (Fig. 1c). Third, more MACs in *S. parvula* increased in abundances contrasting to the depletion of those in *A. thaliana* (Fig. 3d). Amino acids were dominant among *S. parvula* MACs compared to sugars dominant among *A. thaliana* MACs (Fig. 3d). Our results suggest that the two species use two distinct primary metabolite groups to respond to excess K^+^ in addition to contrasting shoot-root responses in line with having only a few MACs under shared responses between the species (Fig. 3b).

We predicted that *S. parvula* was enriched in MACs that minimized cellular stress, while MACs in *A. thaliana* were more representative of pathways disrupted by K^+^ toxicity. Galactose metabolism was significantly enriched among MACs in *S. parvula* roots and *A. thaliana* shoots (Fig. S4a). Galactose metabolism includes several key metabolites (Fig S4b) known to protect plants against oxidative and osmotic stresses (Taji *et al.*, 2002; Nishizawa *et al.*, 2008). Interestingly, these increased uniquely in *S. parvula* roots (Fig. S4b). Antioxidants and osmoprotectants (galactinol, raffinose, myo-inositol, sucrose, and proline) (Taji *et al.*, 2002; Gong *et al.*, 2005; Nishizawa *et al.*, 2008) increased in *S. parvula* compared to *A. thaliana* (Figs 3d, S4b). Collectively, the extremophyte, *S. parvula*, seemed to boost the protective metabolites minimizing damage from oxidative bursts in its roots, while the more stress sensitive species, *A. thaliana*, was depleting its initial pools of such protective metabolites during stress.

### *A. thaliana* transcriptional responses are inept against K^+^ toxicity

We examined the transcriptional response landscapes to deduce distinct cellular processes initiated by *A. thaliana* and *S. parvula* upon excess K^+^. *S. parvula* showed lesser transcriptome-wide variation than *A. thaliana* based on a PCA of 23,281 ortholog pairs (Fig. 4a). Transcriptome similarities between tissues were consistent with high correlations between samples within a tissue/species (Fig. S5). The total number of differentially expressed genes (DEGs) was remarkably higher in *A. thaliana* (9,907 in roots and 12,574 in shoots) compared to those in *S. parvula* (1,377 in roots and 494 in shoots) (Fig. S6 and Table S3).

**Fig. 4.**
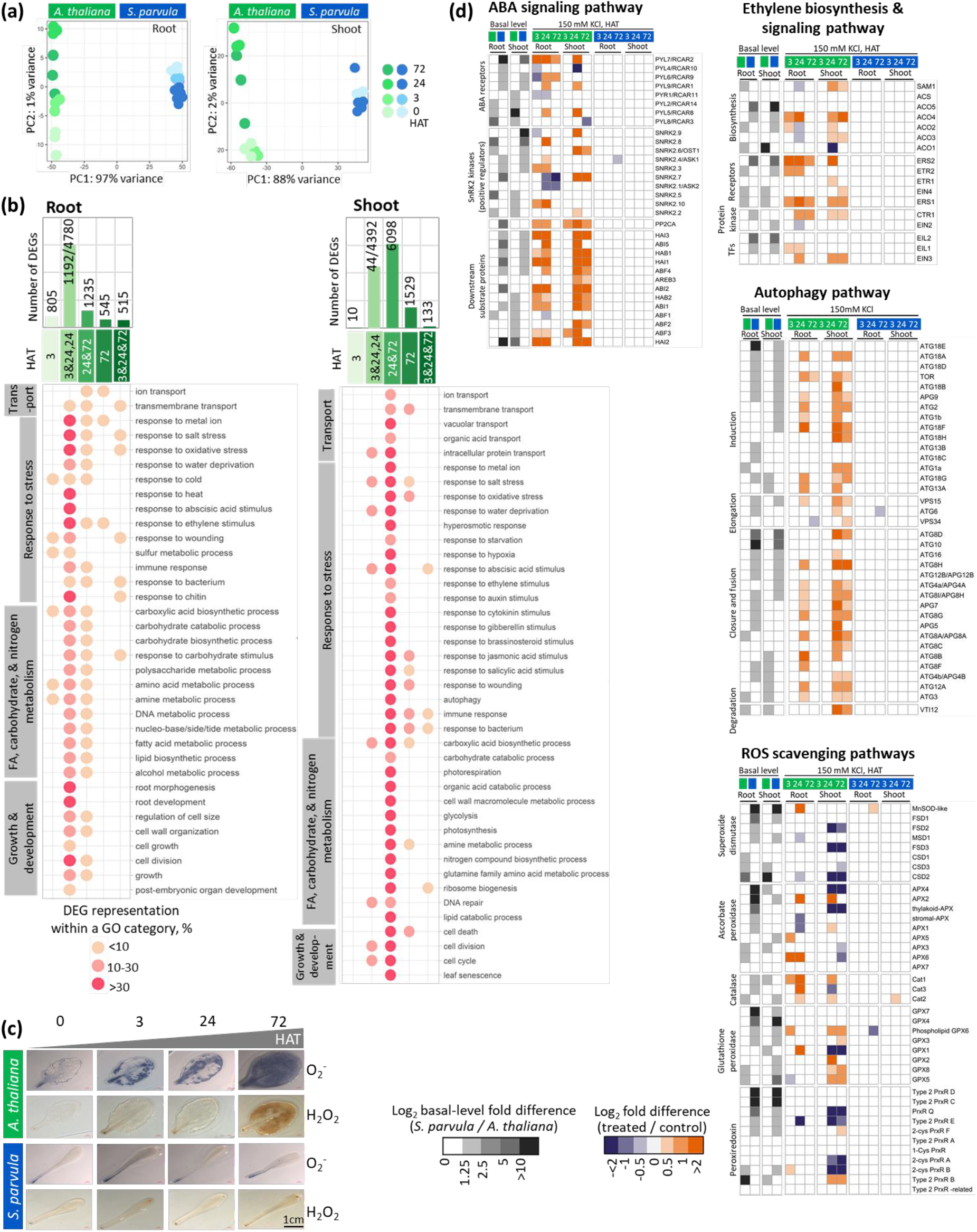
*A. thaliana* shows an overall non-selective transcriptional response during excess K^+^ stress than *S. parvula*. (a) Principal component (PC) analysis of 23,281 ortholog pairs expressed between *A. thaliana* and *S. parvula* root and shoot transcriptomes at 0, 3, 24, and 72 hours after treatment (HAT). (b) Temporally enriched functional processes based on GO annotations associated with differently expressed genes (DEGs) in *A. thaliana*. The temporal sequence is given as 3 h specific, 3 and 24 h shared with 24 h specific, 24 h and 72 h shared, 72 h specific, and present at all-time points from 3-24-72 h. Functional processes that were detected at least in two time points are shown and the processes are sorted based on their functional hierarchy when applicable. (c) Leaves stained for hydrogen peroxide (H_2_O_2_) and superoxide ions (O_2_^-^). Similar growth and treatment conditions used for the RNAseq study. (d) Transcriptional profiles of selected pathways associated with stress signaling. DEGs were called using DESeq2 with a p-adj value based on Benjamini-Hochberg correction for multiple testing set to ≤0.01. Data presented as mean of 3 independent replicates ± SD given using ≥ 5 hydroponically grown plants per replicate.

We annotated all DEGs with a representative Gene Ontology (GO) function (Table S4, S5, S6) and examined their time dependent progression in *A*. *thaliana* (Figs 4b and S6). The temporal transcriptomic response climaxed at 24 and 72 HAT in roots and shoots respectively (Fig. 4b). One of the primary emergent functions enriched in *A. thaliana* was response to stress (Fig. 4b). Within this category, the prevalent specific responses across all time points were responses to salt, oxidative, and ionic stresses. Biotic stress-related GO terms (e.g. response to bacterium, response to chitin, and immune response) were also prominently enriched at all time points (Fig. 4b) in *A. thaliana,* which seemed counterintuitive when *S. parvula* orthologs of those were mostly either unchanged or suppressed (Table S3 and S7). We examined if this was a sign of transcriptional misregulation at the peak response time inept to the extant stress experienced by *A. thaliana*.

We observed three indicators to suggest that there was broad misregulation in transcriptional responses upon excess K^+^ accumulation in *A. thaliana*. First, transcriptional responses were enriched for all major plant hormone pathways in shoots beyond the expected enrichment associated with ABA, suggestive of wide-ranging disruptions to hormone signaling (Fig. 4b). Second, we found autophagy as the most enriched process among DEGs, accompanied by additional enriched processes including cell death and leaf senescence. Third, enriched responses to cold, heat, wounding, drought, and hypoxia suggestive of unmitigated oxidative stress were prevalent during 24-72 HAT (Fig. 4b). Therefore, we assessed the ROS accumulation in *A. thaliana* and *S. parvula* leaves during high K^+^ stress using oxidative stress markers, H_2_O_2_ and O_2_^-^. As predicted by the transcriptional response, leaves of *A. thaliana* showed severe oxidative stress compared to *S. parvula* (Fig. 4c). While fewer DEGs at 72 HAT compared to 24 HAT implied a degree of stress acclimation in *A. thaliana* (Figs 4b and S6), the prolonged ABA signaling together with broad activation of hormone pathways, autophagy, and oxidative stress serve as transcriptional molecular phenotypes to indicate inept responses in *A. thaliana*. These molecular phenotypes become even more compelling when compared to their respective orthologous profiles in *S. parvula* which remained mostly unchanged (Fig. 4d).

Carboxylic acid/carbohydrate and amino acid metabolism formed the second largest group among the most affected processes following stress responses (Fig. 4b). These processes were enriched at all time points in *A. thaliana* roots and shoots consistent with the previous physiological and metabolic responses that showed primary C and N metabolism were severely affected (Figs 1e-f, 2d, 3c-d). Interruptions to photosynthesis at 24 and 72 HAT (Fig. 1e) were aligned with the corresponding transcriptional processes enriched at the same time points in *A. thaliana* shoots (Fig. 4b). Contrarily, root development was detected as a transcriptionally enriched process at 3 HAT (Fig. 4b), although the interruptions to root hair growth was detected at 72 HAT (Fig. 1c).

### *S. parvula* shows targeted transcriptomic responses steered toward stress tolerance

We searched for DEGs among *A. thaliana* whose *S. parvula* orthologs also showed active responses (Fig. 5a). We posited that these orthologs represented cellular processes that require active transcriptional adjustments to survive the accumulation of excess K^+^. We further predicted that the diametric inter-species transcriptional responses (*i.e.* genes that are induced in one species when their orthologs are suppressed in the other species) will be deleterious to the stress sensitive species, while shared responses will be beneficial yet likely underdeveloped or unsustainable to survive prolonged stress in the stress sensitive species (Figs 5a, S6).

**Fig. 5.**
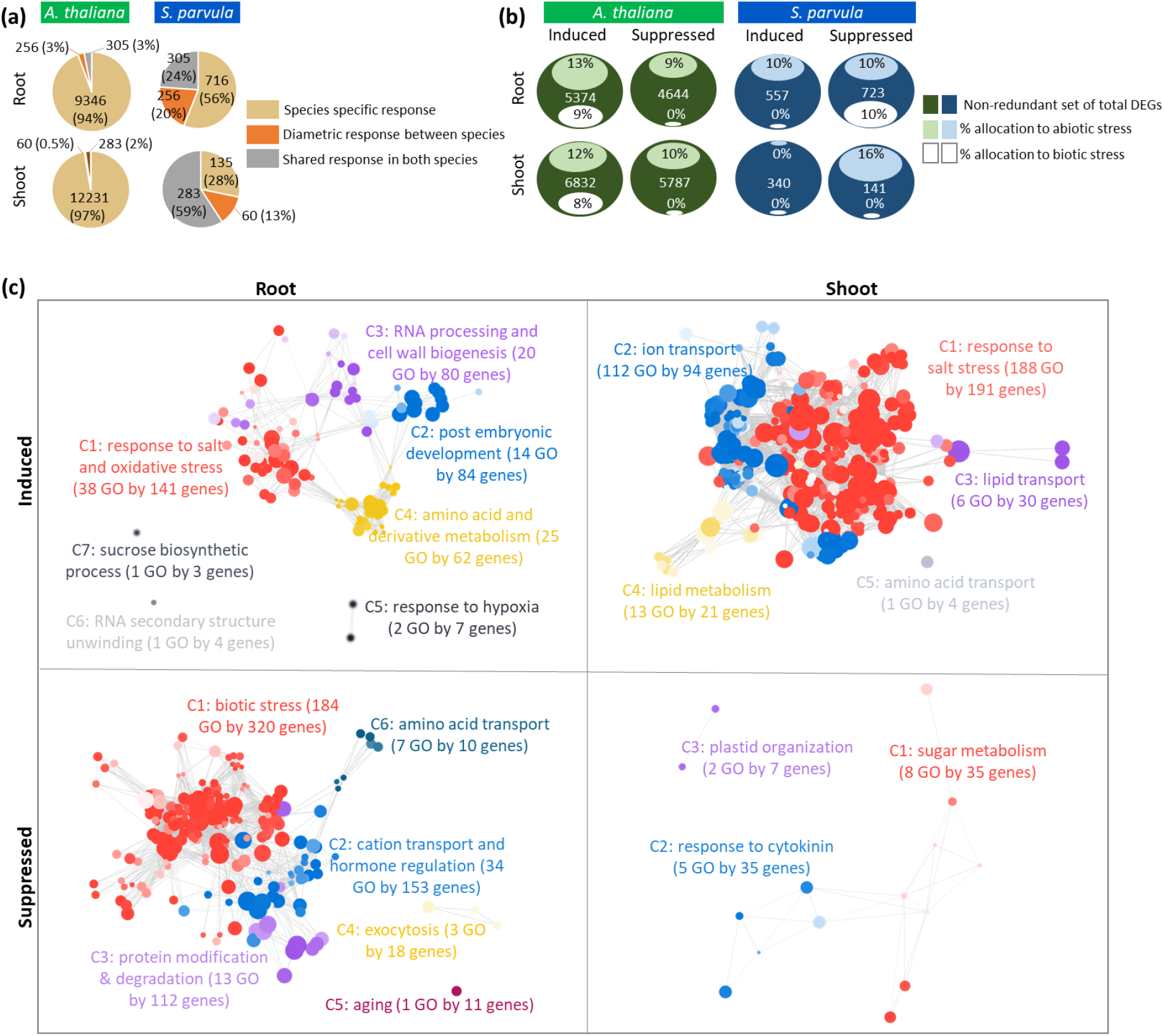
*S. parvula* shows a confined transcriptomic response geared toward concurrent induction of abiotic stress responses and enhanced transcriptional allocation to C and N metabolism. (a) The overall expression specificity and response direction of orthologs in *A. thaliana* and *S. parvula.* Selected orthologs are differentially expressed genes (DEGs) at least in one time point compared to the respective control condition and then counted as a non-redundant set when all 3, 24, and 72 HAT samples were considered for total counts. (b) The proportion contributing to abiotic and biotic stress stimuli within non-redundant DEGs. (c) The functionally enriched processes represented by DEGs in *S. parvula* that responded to high K^+^. A non-redundant set from all time points (3, 24, and 72 HAT) were used. A node in each cluster represents a gene ontology (GO) term; size of a node represents the number of genes included in that GO term; the clusters that represent similar functions share the same color and are given a representative cluster name and ID; and the edges between nodes show the DEGs that are shared between functions. All clusters included in the network have adj p-values ≤0.05 with false discovery rate correction applied. More significant values are represented by darker node colors. DEGs determined at p-adj value set to ≤0.01. Data presented as mean of 3 independent replicates; ± SD given using ≥ 5 hydroponically grown plants per replicate. HAT-Hours after treatment.

Response to stress was the largest functional group representative of diametric responses showing induction in *A. thaliana* compared to suppression in *S. parvula* (Fig. S6). Most of the subprocesses in this cluster were associated with biotic stress (Table S9). Therefore, we further assessed this transcriptional divergence between the two species as a proportional effort invested in biotic vs abiotic stress out of total non-redundant DEGs within each species (Fig. 5b). The effort to suppress biotic stress in *S. parvula* roots (10%) was similar to the proportional induction for biotic stress in *A. thaliana* (9%) (Fig. 5b). Contrarily, orthologs that were suppressed in *A. thaliana*, but induced in *S. parvula* roots were associated with ion transport and cell wall organization (Fig. S6a). These transcriptional adjustments support the physiological response observed for *S. parvula* where uncompromised root growth was coincident to uninterrupted nutrient uptake during exposure to excess K^+^ (Figs 1b, 2c). The orthologs suppressed in *A. thaliana* but induced in *S. parvula* shoots were enriched in carboxylic acid and amine metabolism and transport functions (Fig. S6b). Collectively, the antithetical transcriptomic, metabolic, ionomic, and physiological responses between the two species support the stress resilient growth of *S. parvula* distinct from *A. thaliana* (Figs 1-5).

The orthologs that showed shared inductions (156 in roots and 199 in shoots) were largely represented by abiotic stress responses (Fig. S6 and Table S10). The orthologs with shared suppression (149 in roots and 84 in shoots) were enriched in biotic stress responses in roots and photosynthesis in shoots. However, suppressed orthologs associated with biotic stress in *A. thaliana* (0.004%) were minimal based on a proportional effort compared to that in *S. parvula* (10%) (Fig. 5b).

Over 50% of orthologs differently expressed in response to excess K^+^ in *S. parvula* roots and ∼30% in shoots showed unique expression trends different from *A. thaliana* (Fig. 5a). We postulated that *S. parvula* activates decisive transcriptional regulatory circuits that are either absent (*i.e. S. parvula*-specific responses) or organized differently (*i.e.* diametric responses) than in *A. thaliana* when responding to excess K^+^ stress.

The overall transcriptomic response of *S. parvula* encapsulates induction of more targeted salt stress responses than that of *A. thaliana*, including oxidative stress responses, sugar and amino acid metabolism, and associated ion transport, with concordant induction in growth promoting processes and transcriptional resource recuperation by suppressing biotic stress responses (Fig. 5c). The transcriptional effort to facilitate growth amidst excess K^+^ accumulation in tissues is reflected by induced transcripts involved in cell wall biogenesis, RNA processing, and development along with concurrent suppression for rapid growth limiting processes such as cell wall thickening and callose deposition (Table S10).

### Balance between differential expression of genes encoding K^+^ transporters

The ability in *S. parvula* to curb K^+^ accumulation and prevent the depletion of major nutrients (Fig. 2) led us to hypothesize that genes encoding ion transporters are differentially expressed to prevent nutrient imbalance during excess K^+^ stress. Moreover, ion transport was among the most represented functions within and between species transcriptome comparisons (Figs 4b, 5c, and S6a). We predicted that the genes coding for transporters which allow K^+^ into roots and upload to the xylem or phloem would be primary targets for down-regulation in the effort to restrain K^+^ accumulation, while inducing those that aid in vacuolar sequestration.

We first searched for genes that encoded K^+^ transporters that showed significantly different basal level abundances (Fig. 6a) or/and responses to high K^+^ (Fig. 6b). We categorized those into four transport routes: (a) limit entry into roots, (b) promote efflux from roots, (c) constrain long distance transport between root and shoot, and (d) enhance sequestration into vacuoles. We then assessed routes that were potentially weakened in *A. thaliana* and/or alternatively regulated in *S. parvula*.

**Fig. 6.**
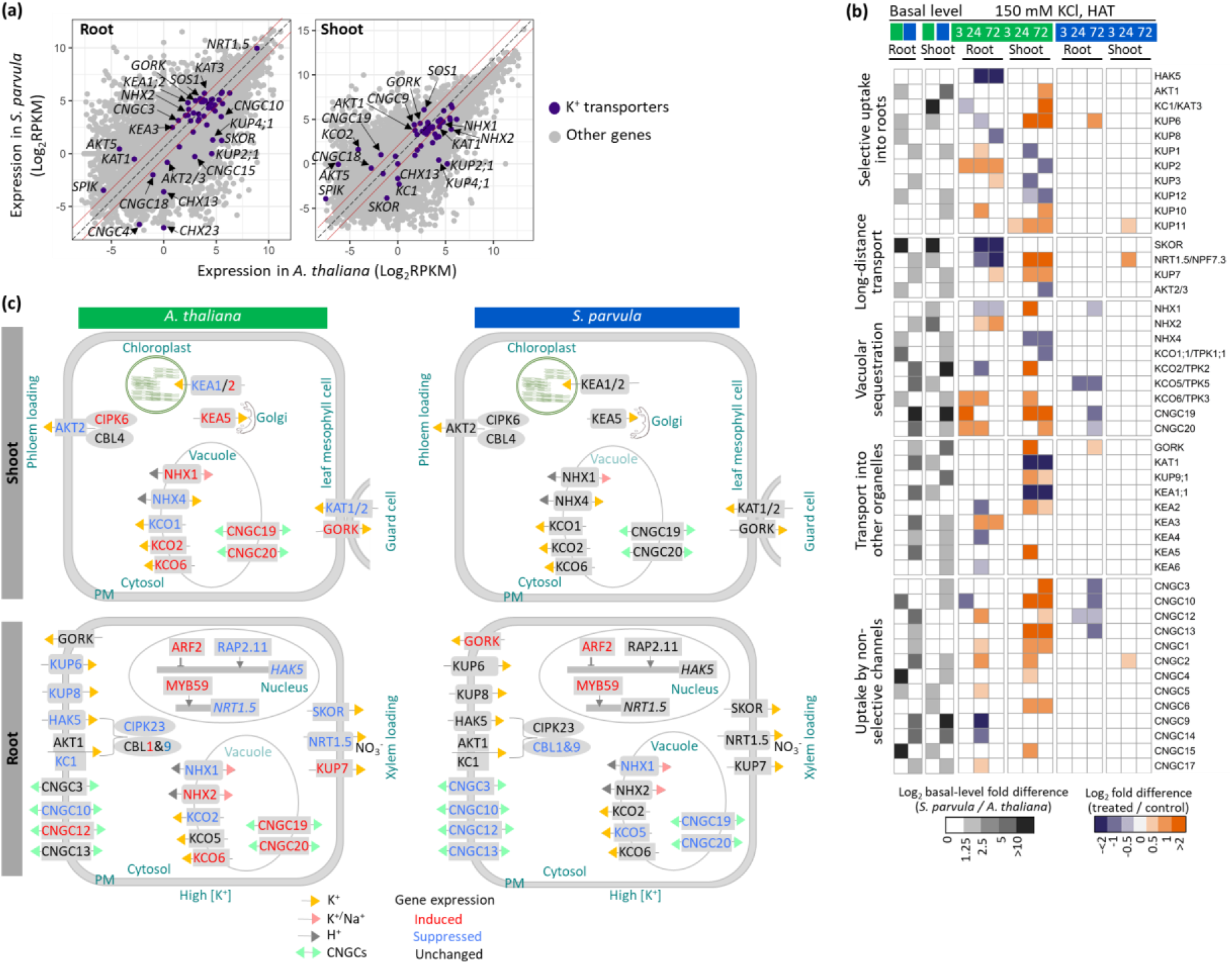
Differential expression of K^+^ transporters in *A. thaliana* and *S. parvula.* (a) Basal level expression comparison of orthologs between *A. thaliana* and *S. parvula* in roots and shoots. The dash-gray diagonal line marks identical expression in both species and solid red lines represent a 2-fold change in one species compared to the other species. Orthologs encoding transporters/channels with ≥2-fold change differences between the species are labeled. (b) The temporal expression profiles of K^+^ transporters and channels in *A. thaliana* and their *S. parvula* orthologs upon high K^+^ treatment. (c) Key K^+^ transporters and channels differently regulated between roots and shoots in *A. thaliana* and *S. parvula*. DEGs determined at p-adj value set to ≤0.01 compared to 0 HAT (Hours after treatment). Data presented as means of 3 independent replicates; ± SD given using ≥ 5 hydroponically grown plants per replicate.

**(a) Limit entry into roots.** The most suppressed gene (13-fold reduction) in *A. thaliana* under excess K^+^ is the *high-affinity K^+^ transporter 5* (*HAK5*) (Gierth *et al.*, 2005), a major transporter for K^+^ uptake during low K^+^ availability (Fig. 6b). Under sufficient K^+^ levels, the low affinity transporter, AKT1 with KC1 is activated (Gierth *et al.*, 2005; Wang *et al.*, 2016). HAK5 and AKT1 are activated by the protein kinase complex, calcineurin B-like proteins 1 and 9 (CBL1&9)/CBL-interacting protein kinase 23 (CIPK23) (Ragel *et al.*, 2015). Under excess K^+^ stress both transporter complexes and their core regulatory unit in *A. thaliana* roots were suppressed (Figs 6b, c and S7a). Interestingly, the orthologous transporters in *S. parvula* roots were unchanged but the orthologs of the interacting kinase complex were suppressed (Figs 6b and S7a). The transcriptional effort to limit entry of K^+^ into roots during K^+^ stress is further exemplified by the suppression of *RAP2.11*, a transcriptional activator of *HAK5* (Kim *et al.*, 2012) and the induction of *ARF2*, a repressor of *HAK5* transcription (Zhao *et al.*, 2016), in *A. thaliana* roots (Fig 6c). Similarly, *ARF2* was induced in *S. parvula* roots. Other K^+^ transporters associated with K^+^ uptake into roots were suppressed in *A. thaliana* (e.g. KUP6 and KUP8) under excess K^+^ stress (Fig 6B). Alternatively, *S. parvula* roots showed a concerted transcriptional suppression of multiple genes encoding cyclic nucleotide gated channels, CNGC3/10/12/13 (Fig. 6b-c). This suggested limitations to non-selective uptake of K^+^ into roots (Gobert *et al.*, 2006; Guo *et al.*, 2008).

**(b) Promote efflux from roots.** K^+^ efflux transporters to specifically extrude excess K^+^ from roots to soil are unknown. However, *S. parvula,* which has evolved in soils naturally high in K^+^, induced transcription of a K^+^ outward rectifying channel, *GORK* (Ivashikina *et al.*, 2001) and the Na^+^ exporter *SOS1* (Shi *et al.*, 2000) in roots (Figs 6b, c and S8). Induction of *GORK* is known to cause K^+^ leakage from roots under biotic and abiotic stresses in plants leading to programmed cell death (Demidchik *et al.*, 2014). We propose that *S. parvula* has evolved to allow export of excess K^+^ via induction of GORK without the destructive downstream consequences of cell death as expected in *A. thaliana* (Figs 6b, c, 4b, and 4d). We also note that the basal expression of *GORK* in *S. parvula* roots is higher than in *A. thaliana* roots (Fig. 6a). SOS1, the antiporter with the highest Na^+^ efflux capacity in roots, is known for its Na^+^ specificity (Oh *et al.*, 2009). Therefore, induction of *SOS1* in *S. parvula* under high K^+^ stress (Fig. S8) is likely an effort to counterbalance the increasing osmotic stress due to elevated K^+^ by exporting available Na^+^ from roots. This explanation fits with Na^+^ being the only ion depleted in *S. parvula* roots during excess K^+^ (Fig. 2c).

**(c) Constrain long distance transport between roots and shoots.** The long-distance transport of K^+^ via xylem loading is mediated by SKOR, NRT1.5, and KUP7 in *A. thaliana* (Gaymard *et al.*, 1998; Han *et al.*, 2016; Li *et al.*, 2017). *SKOR* and *NRT1.5* were suppressed in *A. thaliana* roots as predicted. However, *KUP7* showed induction in *A. thaliana* roots at 72 HAT concordantly when K^+^ accumulation was observed in shoots (Figs 2a and 6b). Contrastingly, none of these transporters were differently regulated in *S. parvula* roots. AKT2 is the dominant channel protein regulating long distance transport via loading and unloading to the phloem (Dreyer *et al.*, 2017). It too is significantly suppressed in *A. thaliana* shoots, but unchanged in *S. parvula* roots and shoots (Fig 6b-c).

**(d) Enhance sequestration into vacuoles.** Vacuolar [K^+^] is spatiotemporally regulated primarily by Na^+^,K^+^/H^+^ antiporters, NHX1 and NHX2 and secondarily with higher selectivity for K^+^ by NHX4 (Bassil *et al.*, 2019). The transcriptional signal to promote K^+^ sequestration in *A. thaliana* roots or shoots is unclear with mixed regulation among *NHXs* compared to a more coordinated co-expression in *S. parvula* shoots (Fig. 6b-c). Furthermore, *A. thaliana* induced *KCO* genes encoding K^+^-selective vacuolar channel known to release K^+^ from vacuoles to the cytosol (Voelker *et al.*, 2006) whereas, *S. parvula*, suppressed *KCO* orthologs implying K^+^ sequestration (Fig. 6b-c). Such an attempt is further reinforced by the suppression of tonoplast localized nonselective cation channels *CNGC19* and *CNGC20* (Yuen & Christopher, 2013) in *S. parvula* roots.

Concordant to decreased photosynthesis, *A. thaliana* shoots represent a molecular phenotype suggestive of closed stomata via an induction of *GORK* together with a suppression of guard cell localized *KAT1/2* (Fig. 6b and c) (Ivashikina *et al.*, 2001; Szyroki *et al.*, 2001). Such a molecular phenotype is absent in *S. parvula*. Additionally, a sweeping array of differentially regulated aquaporins and calcium signaling genes were apparent in *A. thaliana* compared to limited orthologous responses in *S. parvula* (Fig. S7). This reinforces our overall depiction of the stress response in *S. parvula* to reflect a more restrained response during excess K^+^.

### Excess K^+^-induced nitrogen starvation in *A. thaliana* avoided in *S. parvula*

The reduction in total nitrogen and amino acids in *A. thaliana* while those increased in *S. parvula* (Figs 2, 3); followed by suppression of amine metabolism-associated genes in *A. thaliana* when those were induced in *S. parvula* (Figs 4b, 5c, S6b) necessitated further examination on how excess K^+^ may alter N-metabolism in plants. Under low [K^+^_soil_], N uptake in the form of nitrate is tightly coupled to K uptake and translocation within the plant. Many of the K and N transporters or their immediate post-transcriptional activators are co-regulated at the transcriptional level (Coskun *et al.*, 2017). We searched for specific transcriptomic cues to determine how N transport was interrupted under high K^+^, leading to a deficiency in physiological processes needed to maintain growth or creating a shortage of protective metabolites against oxidative and osmotic stress.

The dual affinity nitrate transporter, NRT1.1 (NPF6.3/CHL1) is the main NO_3_^−^ sensor/transporter accounting for up to 80% of NO_3_^−^ uptake from roots (Feng *et al.*, 2020). Within 3 HAT and onwards, *NRT1.1* in *A. thaliana* roots is down-regulated (Fig. 7a). At low [NO_3_^−^ il], NRT1.1 is activated by CIPK23 to function as a high affinity NO_3_^−^ transporter (Coskun *et al.*, 2017). In *A. thaliana* (and not in *S. parvula*) roots, *CIPK23* is concurrently suppressed with the main K-uptake system formed of *HAK5* and *AKT1-KC1* (Fig. 6c). This potentially limits the N content in roots within 24 HAT (Fig. 2b). Correspondingly, *A. thaliana* roots activated N starvation signals by inducing the expression of genes encoding high affinity NO_3_^−^ transporters, NRT2.1 and NRT2.4 (O’Brien *et al.*, 2016) despite sufficient N in the growth medium (Fig. 7A). Therefore, *A. thaliana* showed a molecular phenotype of K^+^-induced N-starvation.

**Fig. 7.**
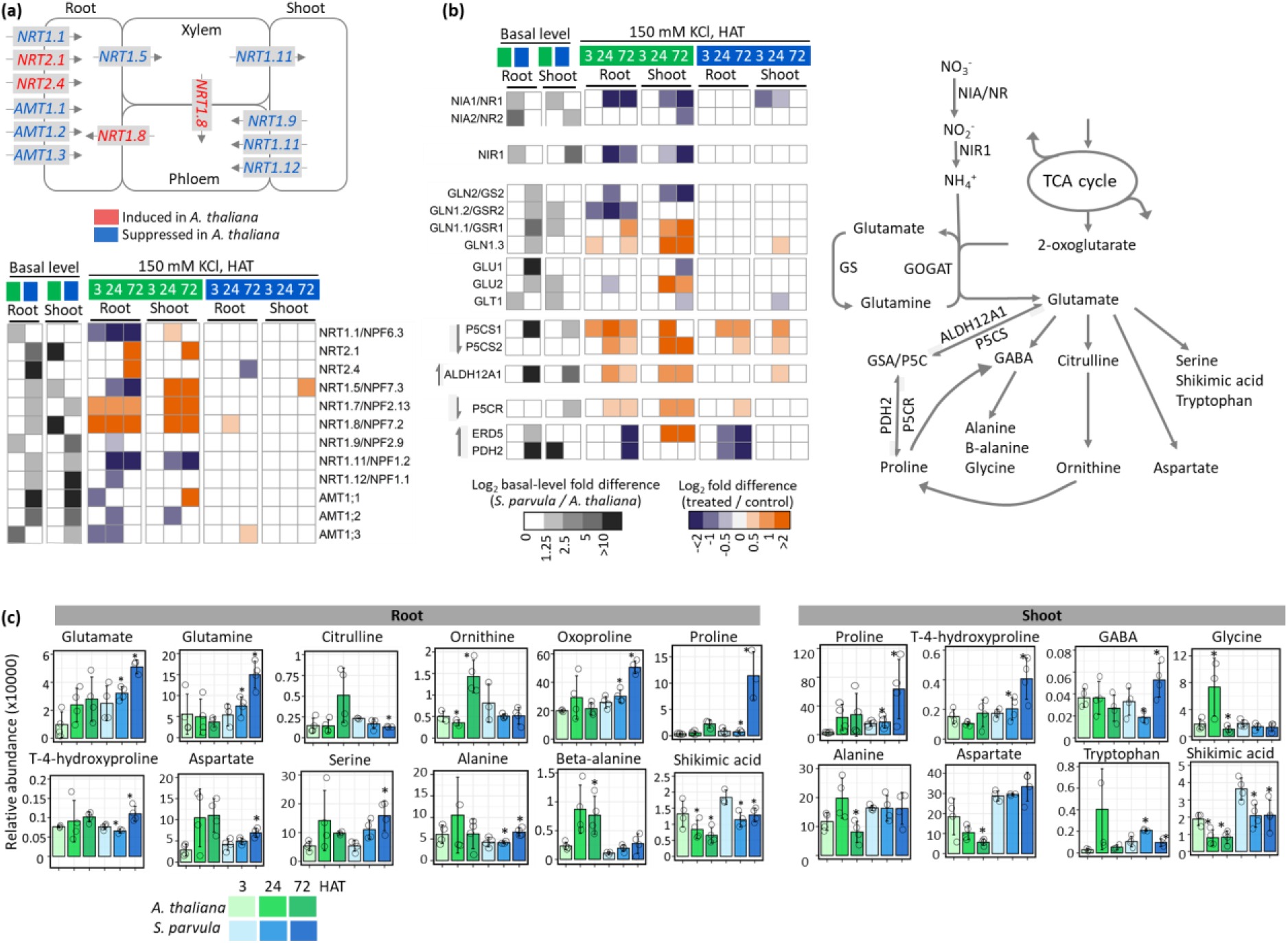
Molecular phenotypes associated with excess K^+^ induced nitrogen starvation and suppressed N-assimilation in *A. thaliana* compared to *S. parvula*. (a) Net expression changes associated with major nitrogen transporters in *A. thaliana* that regulate root uptake and long-distance transport of N. (b) Coordinated transcriptional profiles of nitrogen assimilation and accumulation of glutamate-derived osmoprotectants and antioxidants. The arrows in front of heatmap blocks indicate the direction of the reaction in the pathway. DEGs determined at p-adj value set to ≤0.01compared to 0 HAT (Hours after treatment). (c) Primary metabolites derived from glutamate in roots and shoots. Asterisks indicate significant changes in metabolite abundances based on one-way ANOVA followed by Tukey’s post-hoc test*, p-value* ≤0.05. Data presented as mean of at least 3 independent replicates; ± SD based on ≥ 5 hydroponically grown plants per replicate. Open circles indicate data from each replicate.

The long-distance transport from root to shoot via xylem loading of NO_3_^−^ in roots is primarily regulated via NRT1.5 (NPF7.3) which is a NO_3_^−^/K^+^ cotransporter (Li *et al.*, 2017). In *A. thaliana* (and not in *S. parvula*) roots, *NRT1.5* was suppressed possibly in an attempt to limit excess K^+^ accumulation in shoots, but consequently depriving NO_3_^−^ in shoots (Figs 2b and 7a). High K^+^-induced N-starvation in *A. thaliana* was further reflected by its additional transcriptional effort to remobilize NO_3_^−^ internally. For example, *NRT1.7* and *NRT1.8* were induced to promote translocation of NO_3_^−^ from old to young leaves and from xylem back into roots respectively, while *NRT1.9, NRT1.11,* and *NRT1.12* were suppressed to restrict transport via phloem in shoots (Fig. 7a) (O’Brien *et al.*, 2016). The transcriptional regulatory emphasis on *NRT1.8* is outstanding during excess K^+^, given that it is the highest induced gene (104-fold and 73-fold at 24 and 72 HAT, respectively) in the entire *A. thaliana* transcriptomic response (Fig. 7a and Table S3). Interestingly, *NRT1.8* is triplicated in *S. parvula* (Oh & Dassanayake, 2019), possibly allowing additional regulatory flexibility to redistribute NO_3_^−^ via the xylem back to the roots.

NH_4_^+^ provides another N source and the growth medium included 0.2 mM NH_4_^+^ compared to 1.4 mM NO_3_^−^. Therefore, we expected to see transcriptional induction of NH_4_^+^ transporters to compensate for excess K^+^-induced N-starvation in *A. thaliana* roots. NH_4_^+^ and K^+^ transport are known to be antagonistically regulated (Coskun *et al.*, 2017). The high affinity NH_4_^+^ transporters (AMTs) are inhibited by CIPK23 (Straub *et al.*, 2017). Counterintuitive to our expectations, *AMT1;1/2/3*, which account for >90% of ammonium uptake into roots (Yuan *et al.*, 2007), were co-suppressed in *A. thaliana* (Fig 7A).

We next checked whether the N assimilation pathway from NO_3_^−^ to glutamine via NH_4_^+^ was also suppressed in *A. thaliana*. Indeed, the genes encoding nitrate reductase (*NIA1/NR1, NIA2/NR2*) and nitrite reductase (*NIR*) were coordinately down-regulated in *A. thaliana* under high K^+^ (Fig. 7b). In angiosperms, the main assimilation point of inorganic N to organic compounds is the GS-GOGAT (glutamine synthetase-glutamate synthase) pathway which is tightly coupled to the N and C metabolic state of the plant (O’Brien *et al.*, 2016). The cytosolic *GLN1;2* and plastidial *GLN2* (coding GS enzymes) together with *GLT* and *GLU1* (coding GOGAT enzymes) were suppressed (Figs 2d and 7b). Contrastingly, the induction of *GLN1;1* and *GLN1;3* together with *GLU2* especially in *A. thaliana* shoots may reflect an effort to recycle N under N-starved conditions (Fig. 7b).

We predicted that the suppression of the N-assimilation pathway would be reflected in the change in primary metabolites derived from glutamate in *A. thaliana*. We checked if *A. thaliana* had weakened resources to mount appropriate defenses against osmotic and oxidative stress coincident to the depletion of metabolites directly derived from glutamate that are osmolytes and antioxidants (Fig. 7c). Both species showed a coordinated effort to accumulate proline and its immediate precursors via induction of key proline biosynthesis genes (Fig. 7b, *P5CS1/2*, *P5CR*). However only *S. parvula* was able to significantly accumulate proline during exposure to excess K^+^ (Fig. 7c). Proline has dual functions as an osmoprotectant and an antioxidant (Hayat *et al.*, 2012). We see similar pronounced efforts in increasing antioxidant capacity via GABA and beta-alanine (Fig. 7b, c), concordant to increased synthesis of raffinose and myo-inositol against osmotic stress in *S. parvula* (Fig. S4b). Overall, *S. parvula* is able to accumulate carbon and nitrogen-rich antioxidants and osmoprotectants by maintaining N uptake from roots and N-assimilation pathways independently from the suppressed K-uptake pathways. Contrastingly, the two processes were jointly suppressed in *A. thaliana* leading to the depleted N resources (Fig. 2b) and, in turn, failure to accumulate C and N-rich protective metabolites (Fig. 7c).

### Co-expressed gene clusters indicate stress preparedness in *S. parvula*

We generated co-expressed clusters using 14,318 root and 14,903 shoot ortholog pairs of which, we identified five root and three shoot clusters (Fig. 8, RC1-5 and SC1-3, respectively) (Fig 8. and Table S11). In three root co-expressed clusters, *A. thaliana* orthologs showed a maximum response at 24 HAT, while *S. parvula* showed constitutive responses (Fig. 8a, RC1-3). These clusters largely represented transcripts associated with stress responses, C and N metabolism, transport, and root development we discussed earlier (Fig. 4). The 4th and 5th clusters (Fig. 8a, RC4-5, 203 ortholog pairs), where *S. parvula* showed a response, comprised functionally uncharacterized genes (37%) that could not be summarized into representative processes. This highlights the extent of functional obscurity or novelty of genes that respond to specific ionic stresses minimally characterized in *A. thaliana* (Fig. 8a, RC5), and the novel regulatory modes detected in orthologs of closely related species whose functional assignment may have been overlooked due to the lack of responses in *A. thaliana* (Fig. 8a, RC4). In all three co-expressed shoot clusters, *A. thaliana* again showed a peak response at 24 HAT, while *S. parvula* orthologs showed constitutive expression (Fig. 8b). The enriched functions in shoot clusters largely overlapped to include stress responses and C and N metabolic processes.

**Fig. 8.**
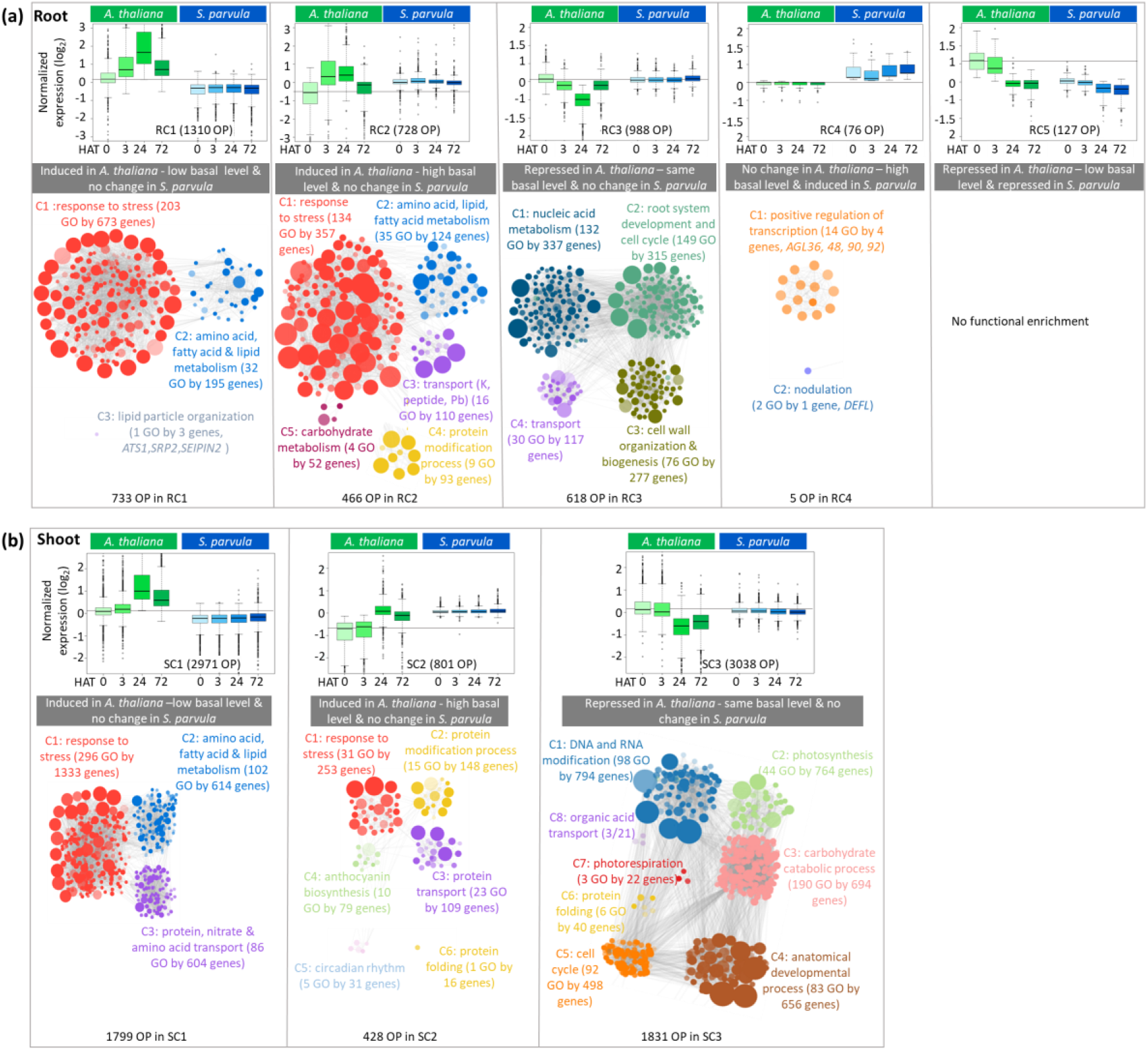
Co-expressed ortholog gene modules highlight stress preparedness for excess K^+^ in *S. parvula* orthologs that are constitutively expressed compared to induction or suppression of *A. thaliana* orthologs. Normalized gene expression clusters of ortholog pairs (OP) between *A. thaliana* and *S. parvula* in (a) roots and (b) shoots. Fuzzy K-means clustering of temporally co-expressed OPs with a membership cutoff of ≤0.5. Box and whisker plots present the median expression at each time point given by the thick line within the box; interquartile range between first and third quartile given by the height of the box; and interquartile range x 1.5 marked by whiskers for lower and upper extremes. Basal level taken as 0 HAT (Hours after treatment) in *A. thaliana* is marked by a grey line in all plots. Each cluster was used for a functional enrichment analysis represented by GOMCL summaries placed below co-expression plots. A node in each cluster represents a gene ontology (GO) term; size of a node represents the number of genes included in that GO term; the clusters that represent similar functions share the same color and are given a representative cluster name; and the edges between nodes show the orthologs that are shared between functions. All clusters included in the network have adj p-values ≤0.05 with false discovery rate corrections applied. More significant enrichments are represented by darker nodes.

*S. parvula* showed constitutive expression in all clusters (9,633 orthologs) except in RC4 and RC5, while *A. thaliana* showed constitutive expression only in RC4 (76 orthologs) (Fig. 8). Overall, these co-expressed clusters between *A. thaliana* and *S. parvula* demonstrate the transcriptome-level stress preparedness in *S. parvula* to facilitate growth and development during excess K^+^ stress.

## Discussion

Salt tolerance mechanisms against high K^+^ are largely unknown compared to the collective understanding for high Na^+^ tolerance in plants. Our results demonstrate that high K^+^ is more deleterious than Na^+^ given at the same external concentrations (Fig. 1). Previous studies support this observation noting that excess KCl caused more severe stress symptoms (Eijk, 1939; Ashby & Beadle, 1957; Eshel, 1985; Matoh *et al.*, 1986; Cramer *et al.*, 1990; Wang *et al.*, 2001; Ramos *et al.*, 2004; Richter *et al.*, 2019; Zhao *et al.*, 2020). The canonical adaptations described for salt tolerance mechanisms associated with NaCl-induced salt stress (Pantha & Dassanayake, 2020) are insufficient to explain adaptations required for KCl-induced salt stress.

The extremophyte, *S. parvula,* amidst high K^+^ can sustain its growth and development; compartmentalize excess K in roots than in shoots; maintain uninterrupted nutrient uptake; increase its antioxidants and osmoprotectants; decouple transcriptional regulation between K and N transport; and coordinately induce abiotic stress response pathways along with growth promoting pathways (Fig. 9). Contrastingly, the more stress-sensitive plant, *A. thaliana* shows, interrupted growth; excessive accumulation of K^+^ in roots and shoots; depletion of essential nutrients; depletion of N-containing metabolites; and sweeping transcriptomic adjustments suggesting initiation of autophagy, ROS accumulation, induction of both abiotic and biotic stresses, and responses to all major hormone pathways (Fig. 9). Based on our comparative analyses, we propose two deterministic steps in the overall stress response sequence to survive high K^+^ stress.

**Fig. 9.**
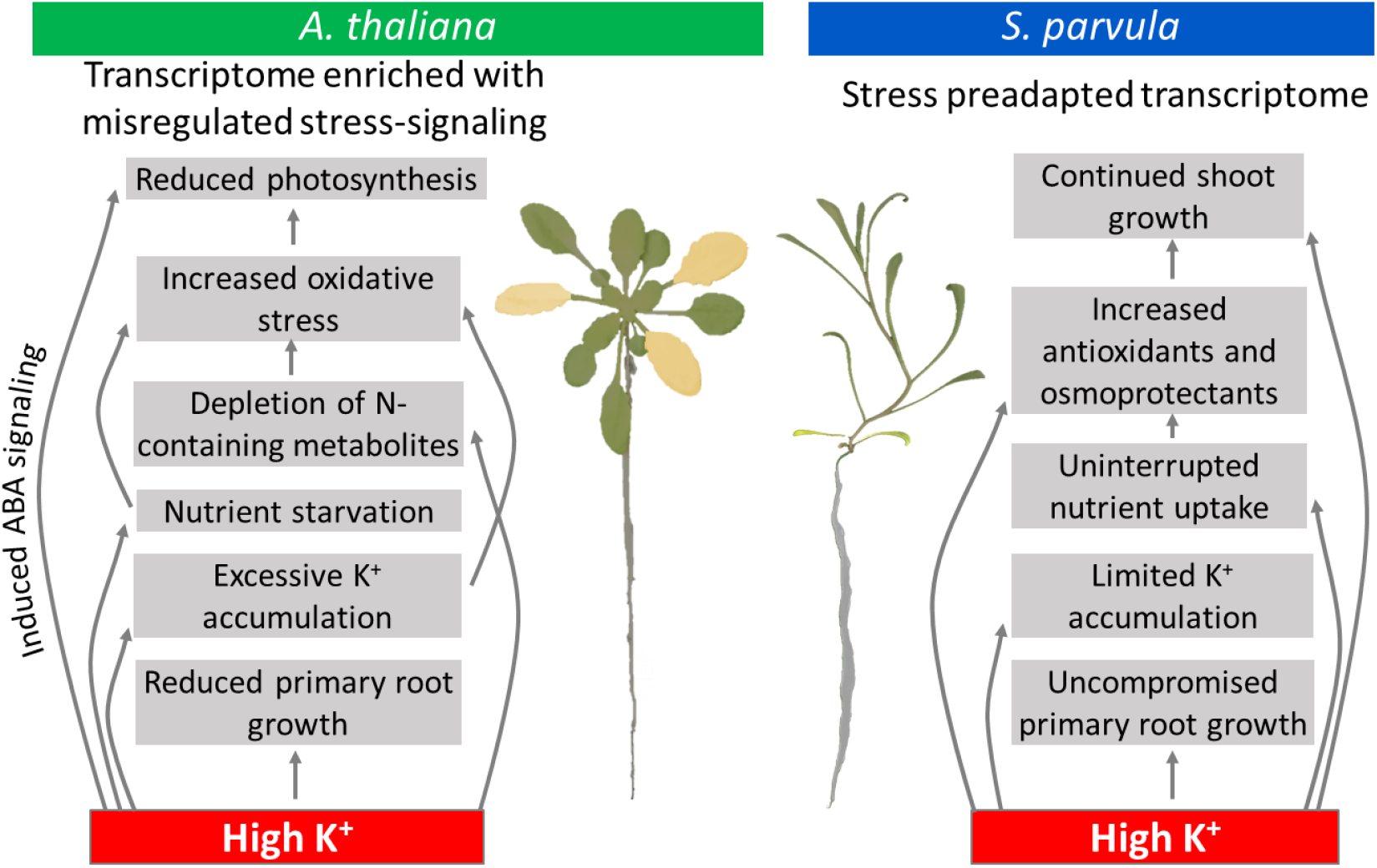
Cellular processes that determine stress resilient growth from stress-affected growth during K-induced salt stress.

### Surviving K toxicity by avoiding N starvation

K is a macronutrient and plants have evolved many functionally redundant transporters to uptake K^+^ into roots and redistribute within plants (Shabala & Cuin, 2008). When external [K^+^] exceed physiologically optimal conditions, it is not surprising that the immediate response from both plants was to suppress expression of K^+^ transporters that primarily control K^+^ influx at the root-soil interface (Fig. 6). Additionally, *S. parvula* down-regulated non-selective CNGCs that may be permeable to K^+^ in roots. Several CNGCs are reported to allow Na^+^ or K^+^ transport and have been implicated in their functions during Na-induced salt stress by controlling Na influx into roots. However, their functional and spatiotemporal specificity remains largely unresolved (Dietrich *et al.*, 2020) and needs to be determined before evaluating how selected CNGCs may be involved in limiting excess K influx under K-induced salt stress.

*A. thaliana* further seems to suppress long distance transport of K^+^ via NRT1.5 that co-transports NO_3_^-^ and K^+^ (Li *et al.*, 2017). NO_3_^-^ is transported as a counterion with K^+^ in root to shoot translocation as described by the ‘Dijkshoorn–Ben Zioni model’ (Dijkshoorn *et al.*, 1968; Zioni *et al.*, 1971; Coskun *et al.*, 2017). The suppression of *NRT1.5* limits NO_3_^-^ remobilization in plants (Chen *et al.*, 2012). This interference to N transport within the plant is compounded by the transcriptional co-suppression of NRT and AMT transporters known to limit N intake from soil (Fig. 7) (Tegeder & Masclaux-Daubresse, 2018). This creates an N-starved condition for *A. thaliana* not observed for *S. parvula.* During limiting K^+^ conditions, N-uptake is down regulated to prevent excess N-induced toxicity in plants as a favorable mechanism to adapt to K^+^-starvation (Armengaud *et al.*, 2004). This interdependent N and K transport and regulation favorable at low [K^+^_soil_] appear to be detrimental at high [K^+^] as it creates an antagonistic pleiotropic effect (condition dependent traits that can cause positive as well as negative impacts).

We observed a significant induction of *NRT1.8/NPF7.2* suggestive of an effort to reimport N from the stele in *A. thaliana* (Li *et al.*, 2010). However, this transcriptional effort did not cascade to the ionomic level (Fig. 2b and d). N remobilization via induction of *NRT1.8* while concurrently suppressing *NRT1.5* (Fig. 7a) during N starvation is regulated by ethylene-jasmonic acid signaling together with low N-sensing by nitrate reductase (Chen *et al.*, 2012) (Zhang *et al.*, 2014). Both ethylene and jasmonic acid signaling are among the enriched differently regulated transcriptional processes in *A. thaliana* (Fig. 4b). Interestingly, *S. parvula* appears to have a more flexible and effective regulatory capacity to allow N-uptake decoupled from restricted K-uptake and it does not suppress internal remobilization of NO_3_^-^ and K^+^ via NRT1.5 (Figs 6 and 7). This may prevent *S. parvula* from experiencing a high-K induced N-starvation.

The depletion of N uptake in *A. thaliana* further cascades into depletion of primary metabolites containing N (Fig. 3d) with a concomitant transcriptional suppression observed in N assimilation via the GS-GOGAT pathway (Fig. 7) (O’Brien *et al.*, 2016; Ji *et al.*, 2019). This not only creates a shortage of essential primary metabolites required for growth and development, but also depletes essential antioxidants and osmolytes to defend against the mounting oxidative and osmotic stresses (Figs 3, 4b-d, 7, 9). High K in the growth medium is known to exert osmotic and oxidative stress (Osakabe *et al.*, 2013; Zheng *et al.*, 2013). This creates an overall need to boost osmotic and antioxidant defense systems to successfully survive high K^+^ toxicity.

Synergistic transcriptional and metabolic resource allocation to increase osmoprotectants during high K^+^ is much more pronounced in *S. parvula* than in *A. thaliana* (Figs 3, 7). Proline accumulation resulting from increased synthesis and reduced catabolism have been widely shown as a key adaptation during salt stress (Kishor *et al.*, 1995; Gong *et al.*, 2005; Kant *et al.*, 2006; Kumar *et al.*, 2010; Hayat *et al.*, 2012).

K and N regulate phosphorus uptake (Coskun *et al.*, 2017; Maeda *et al.*, 2018; Cui *et al.*, 2019), while K^+^ toxicity can induce P-starvation (Ródenas *et al.*, 2019). *A. thaliana* experienced severe shortages of multiple key nutrient depletions (Fig. 2), which *S. parvula* seemingly avoided by having independent regulatory capacity of K and N uptake (Figs 6, 7, and 9). Therefore, we propose that the ability to regulate independent K^+^ uptake is the first key deterministic step towards building resilience to excess K^+^.

### Avoiding transcriptional misregulation of K^+^ signaling

Multiple hormonal pathways use K^+^ for developmental, biotic stress, and abiotic stress signaling (Zhang *et al.*, 2014; Hauser *et al.*, 2017; Shabala, 2017). Canonical mechanisms involving K^+^ signaling for growth are based on sensing external K^+^ at low or favorable conditions. When supplied with toxic levels of K^+^, *A. thaliana* induced non-selective hormone signaling pathways inept to the extant stress (Fig. 4) indicative of transcriptional misregulation. Therefore, we propose that the capacity to avoid transcriptional misregulation of K^+^ signaling is the second major deterministic step in surviving high K^+^ stress. If unavoided, it can lead to systemic damage via activation of ROS and autophagy pathways, as demonstrated by *A. thaliana* with its increased ROS accumulation perhaps resulting from induction of futile biotic stress responses or unmitigated oxidative stresses induced by high K^+^ detrimental especially at a nutrient starved environment (Fig. 4). ROS accumulation combined with autophagy are associated with abiotic and biotic stress responses, and developmental processes (Liu *et al.*, 2005; Thompson *et al.*, 2005; Lv *et al.*, 2014; Pantha & Dassanayake, 2020). However, uncontrolled initiation of autophagy signifies a failed stress response strategy (Floyd *et al.*, 2015). In *A. thaliana* shoots, autophagy is the most enriched transcriptional pathway. The collective transcriptional signal enriched for lipid catabolism, protein degradation, DNA repair, cell death, and leaf senescence (Das & Roychoudhury, 2014) (Fig. 4b) further indicates the maladaptive stress response shown by *A. thaliana* contrasted against a pre-adapted state observed for *S. parvula*, when exposed to excess K^+^ (Figs 5 and 8). Previous transcriptome and metabolome characterizations from extremophytes including *S. parvula* have shown similar stress-ready states for other abiotic stresses (Kant *et al.*, 2006; Lugan *et al.*, 2010; Oh *et al.*, 2014; Wang *et al.*, 2021).

In conclusion, upon exposure to high K^+^, plants undergo physiological, metabolic, and transcriptional changes and a subset of those changes lead to stress adaptive traits while the other responses are indicative of failed cellular responses unable to meet the increasing systemic toxicity exerted by excess K^+^ accumulation. The deterministic steps whether a plant would be able to survive K-induced salt stress or descend into unmitigated stress responses were primarily dictated by the ability to regulate K uptake independent from other nutrient uptake pathways while avoiding deleterious signaling processes. This decoupled regulation of K transport and stress signaling can be targeted to design improved crops that are better able to dynamically adjust to a wide array of soils or irrigation water sources with different salt compositions increasingly comprised of high K in water-limited environments.

## Acknowledgements

We thank Drs. Guannan Wang, Kieu-Nga Tran, Aaron Smith, and John Larkin, and Chathura Wijesinghege for providing feedback on the manuscript and facilitating helpful discussions; Prava Adhikari for additional assistance with phenotyping, and undergraduate students Saad Chaudhari, Megan Guilbeau, and August Steinkamp for their assistance to grow plants. This work was supported by the US National Science Foundation awards MCB-1616827 and IOS-EDGE-1923589, US Department of Energy BER-DE-SC0020358, and the Next-Generation BioGreen21 Program of Republic of Korea (PJ01317301) awarded to MD and DHO. PP was supported by an Economic Development Assistantship award from Louisiana State University. We acknowledge LSU High Performance Computing services for providing computational resources, and Drs. John Cheeseman and Alvaro Hernandez for their assistance with sequencing services at the University of Illinois.

## Author Contributions

PP conducted experiments and performed data analyses. DL supervised and assisted with measuring CO_2_ assimilation rates. DHO provided bioinformatics assistance. MD developed the experimental design and supervised the overall project. PP and MD interpreted results and wrote the article with input from all co-authors who revised and approved the final manuscript.

## Data Availability

The RNA-seq reads generated in this study are available at NCBI-SRA database under BioProject PRJNA63667. Mapped transcripts from this study for *S. parvula* can be browsed at https://www.lsugenomics.org/genome-browser.

## Supplementary figure caption

**Fig. S1.**
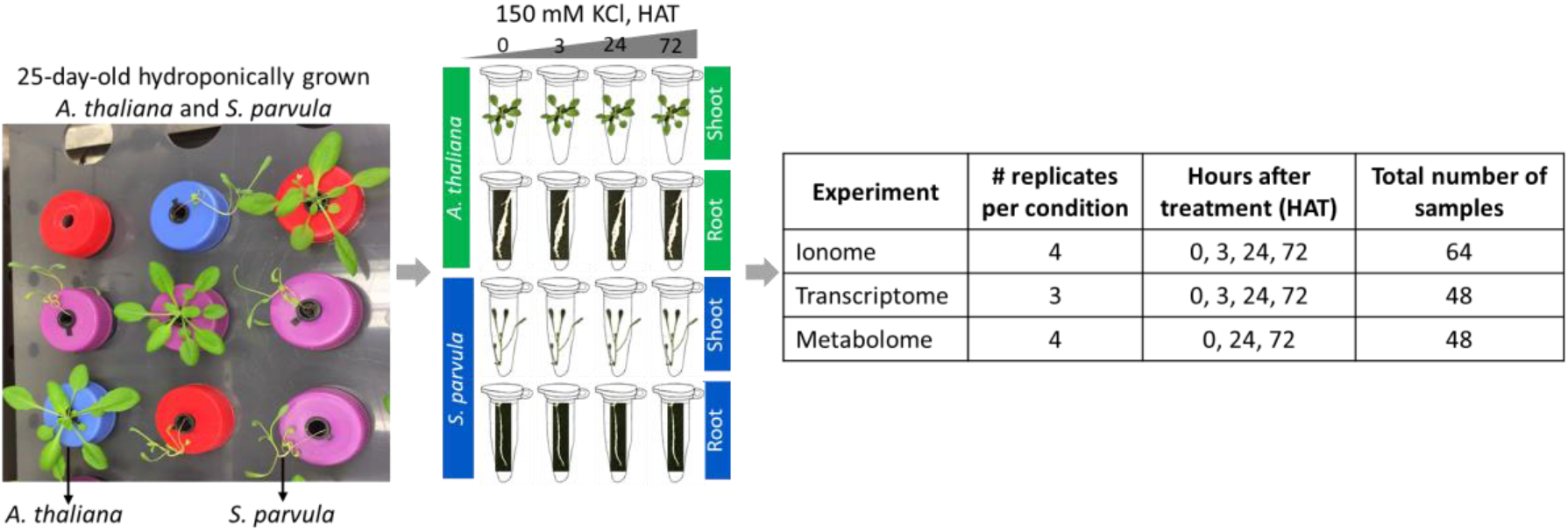
Sampling scheme for the ionome, metabolome, and transcriptome assessements performed in this study. 25-day-old *Arabidopsis thaliana* and *Schrenkiella parvula* plants were grown in 1/5^th^ Hoagland’s solution with/without supplemental 150 mM KCl for up to 72 hours after treatment (HAT). All samples were 28-day-old at the time of harvest. Shoot and root tissue samples were harvested on a randomized basis from the same growth chamber, at the same time of day for control and salt-treated plants. Roots were briefly dried with a paper towel to soak any excess growth solution. All treatment and harvest times were set at 4 h after the beginning of the light cycle to avoid variation due to circadian effects. The salt treatment was non-lethal to both *A. thaliana* and *S. parvula* plants based on preliminary tests using a series of salt concentrations. Both ionome and metabolome profiles have 4 biological replicates and transcriptome samples have 3 biological replicates. Each replicate contains tissues from at least 5 different plants.

**Fig. S2.**
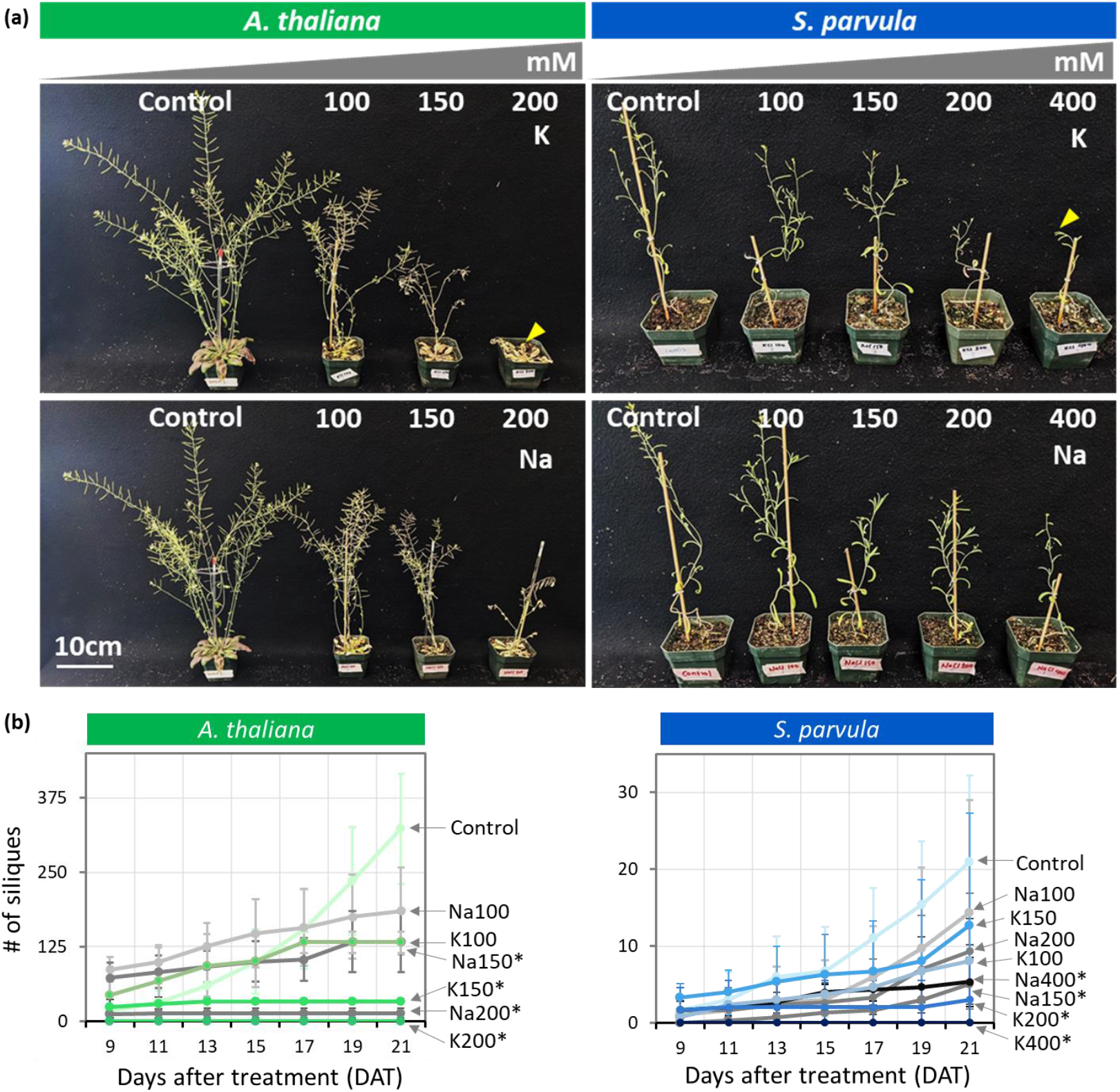
KCl is more toxic than NaCl at the same osmotic strength at the end of the lifecycle. (a) 21-day-old plants treated for 14 days (salt applied every other day). *A. thaliana* plants treated with KCl show severe growth disruptions compared to the treatments given with the same concentration of NaCl. In *A. thaliana*, 200 mM KCl treated plants did not develop any flowers at the completion of the experiment with 20 days of monitoring after treatment ended, whereas plants treated with 200 mM NaCl flowered and developed siliques. *S. parvula* showed a similar observation at 400 mM KCl treatment compared to NaCl treatment. Yellow arrowhead indicates the absence of floral development. (b) Number of siliques quantified from NaCl and KCl treated plants shown in (a). A minimum of 3 plants per condition were used for both species. Asterisks indicate significant changes between the treated samples to its respective control samples (t-test with p ≤0.05). Data presented as mean of at least 3 independent biological replicates with ± SD. DAT-Days after treatment.

**Fig. S3.**
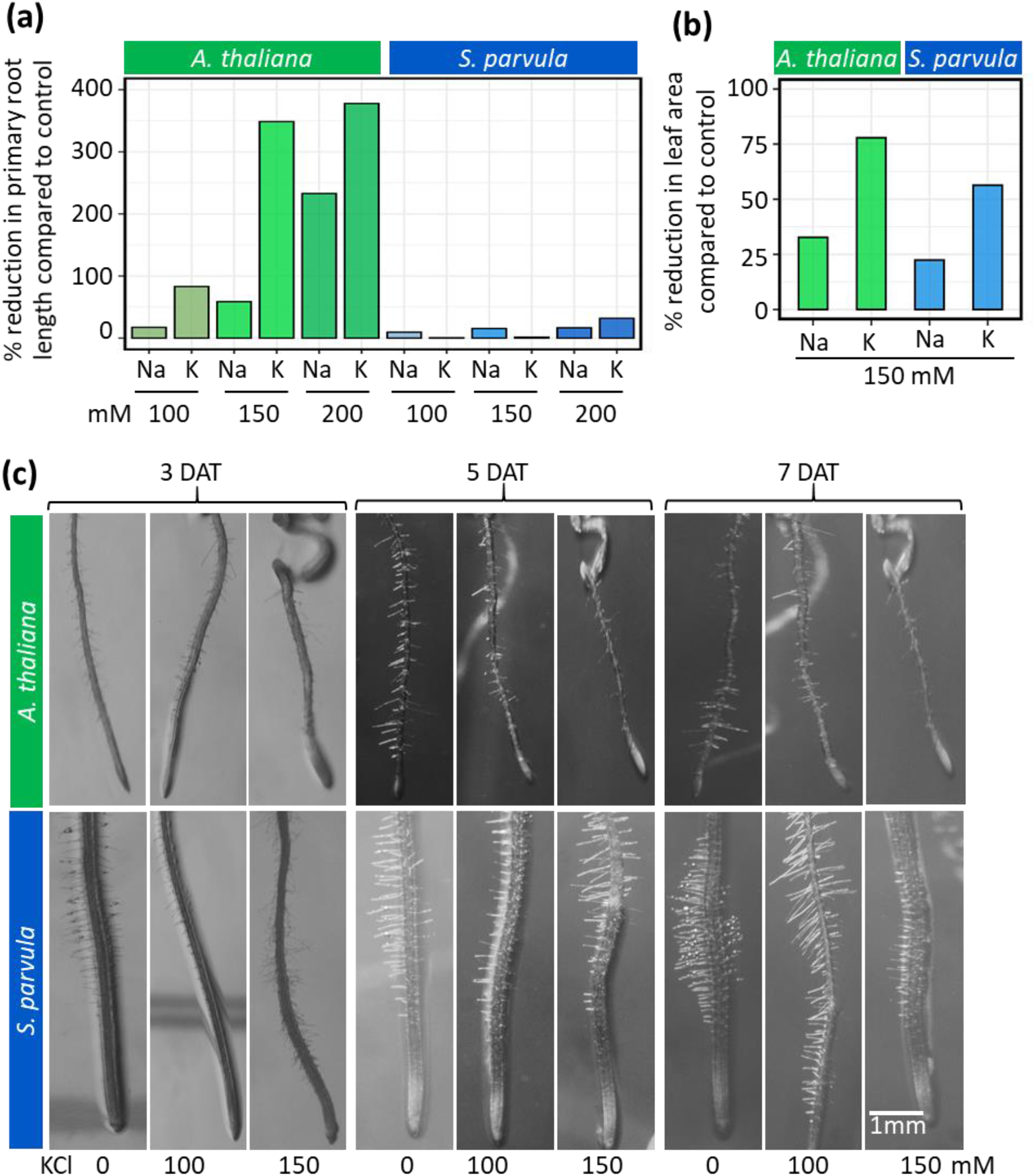
KCl is more toxic than NaCl at the same osmotic strength at the vegetative growth phases. Effects on (a) primary root length, (b) leaf area, and (c) root hair development upon high Na and K treatments compared to their respective controls. Primary root length was measured on 17-day-old seedlings grown on 1/4^th^ MS media supplemented with 0 to 200 mM NaCl or KCl based on a 12-day treatment. Total leaf area per plant measured when plants developed the first floral bud following 1-2 weeks of salt treatments in a hydroponic medium. Minimum 5 plants per condition used in (a) and a minimum of 3 plants per condition used for (b). Same roots were photographed on 3, 5, and 7 DAT (Days after treatment).

**Fig. S4.**
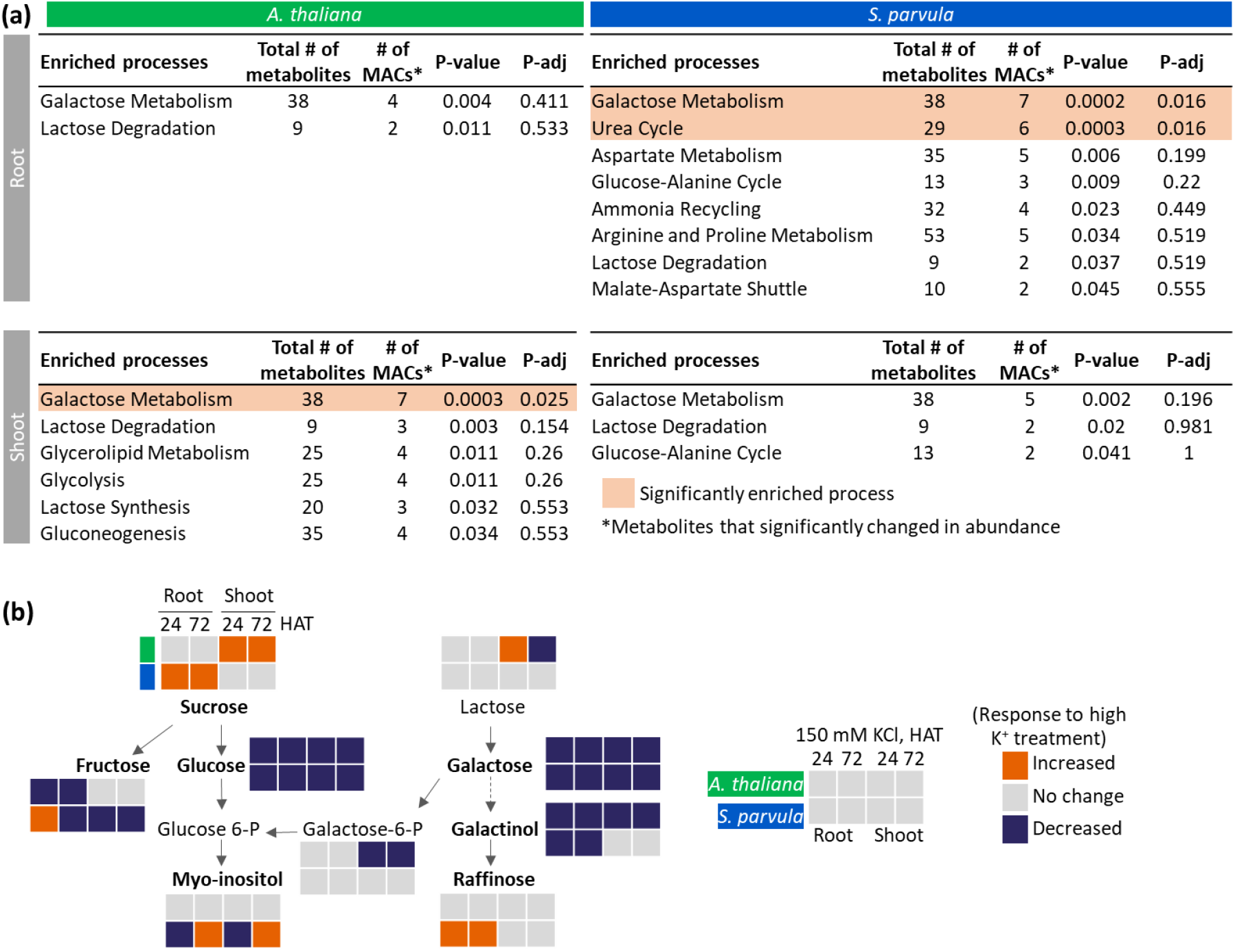
Carbohydrate and nitrogen metabolism related processes were enriched in *S. parvula* roots. (a) Enriched metabolic pathways in *A. thaliana* and *S. parvula* in response to excess K stress. Metabolite Enrichment Analysis was performed with MetaboAnalyst with a p-adj cut off set to ≤0.05. p-values were adjusted using Benjamini-Hochberg correction for multiple testing. (b) Simplified galactose metabolism pathway (KEGG Pathway ID:ath00052). The seven metabolites that significantly changed in abundance and quantified using GC-MS are indicated in bold font.

**Fig. S5.**
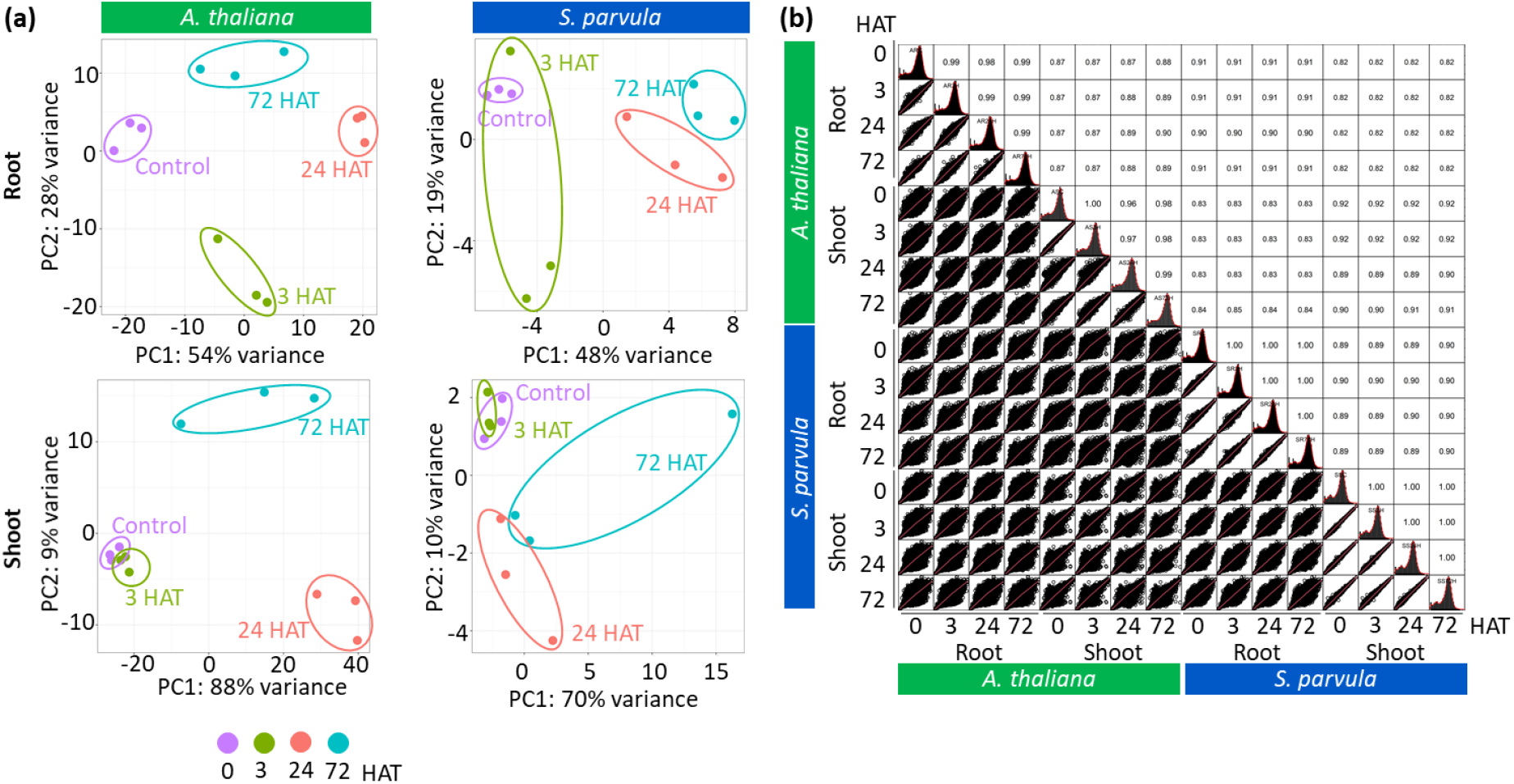
Overview of transcriptomes in each condition tested in response to high K stress in *A. thaliana* and *S. parvula* roots. (a) Principal component analysis (PCA) for 0, 3, 24, and 72 hours after treatment (HAT) for *A. thaliana* and *S. parvula* roots and shoots. (b) Overall transcript level correlation between conditions. Correlation plots were generated using PerformanceAnalytics library in R 4.0.2. Pearson correlation coefficient is given for each comparison.

**Fig. S6.**
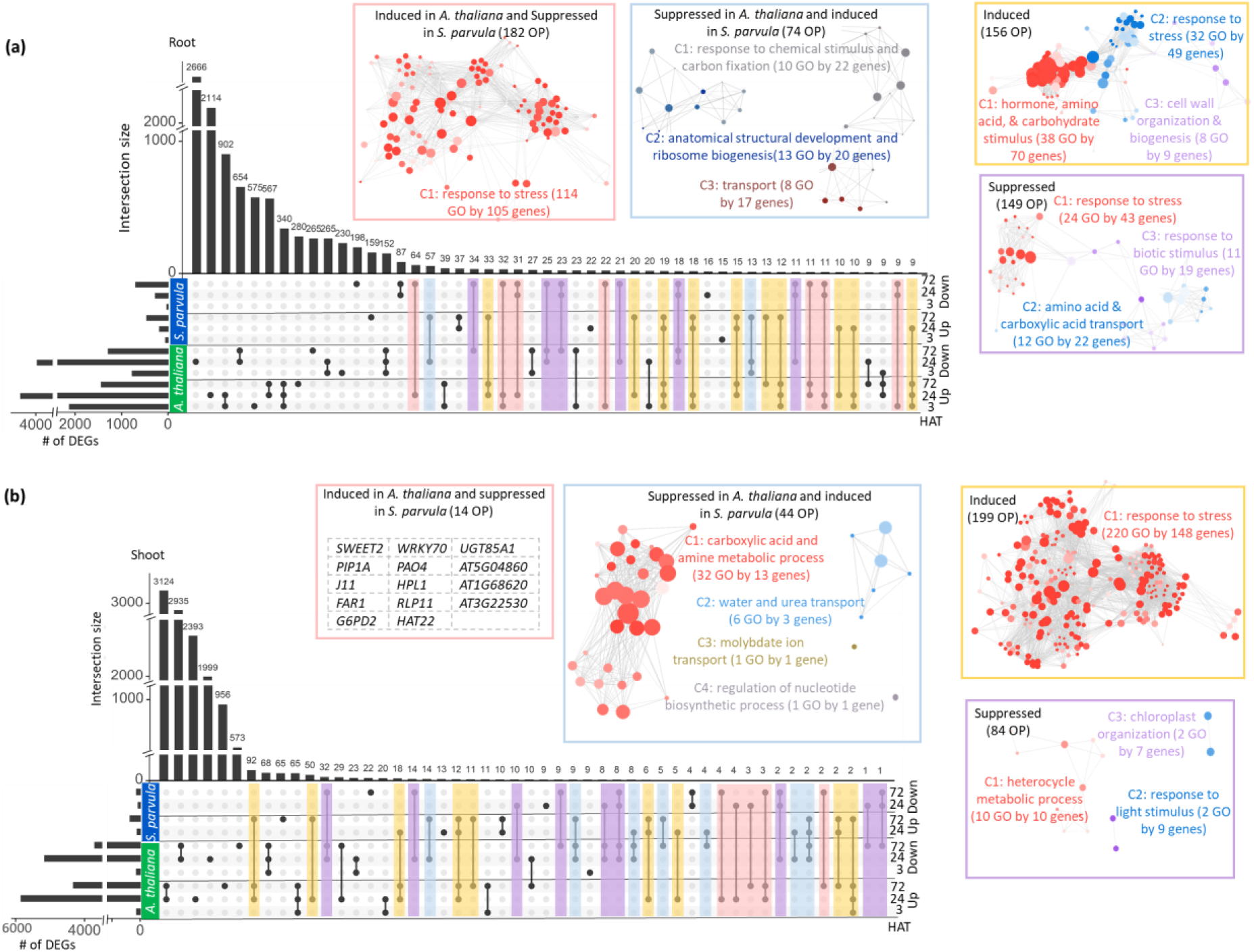
The majority of differentially expressed genes (DEGs) show species specific responses followed by tissue and response time specificity. Differentially expressed genes (DEGs) between *A. thaliana* and *S. parvula* for (a) roots and (b) shoots. Enriched functions are highlighted for diametric and shared responses between differently regulated ortholog pairs (OP). DEGs at each time point were called using DESeq2 compared to 0 h with a p-adj value based on Benjamini-Hochberg correction for multiple testing set to ≤0.01. The shared genes were plotted using UpsetR in R 4.0.2.

**Fig. S7.**
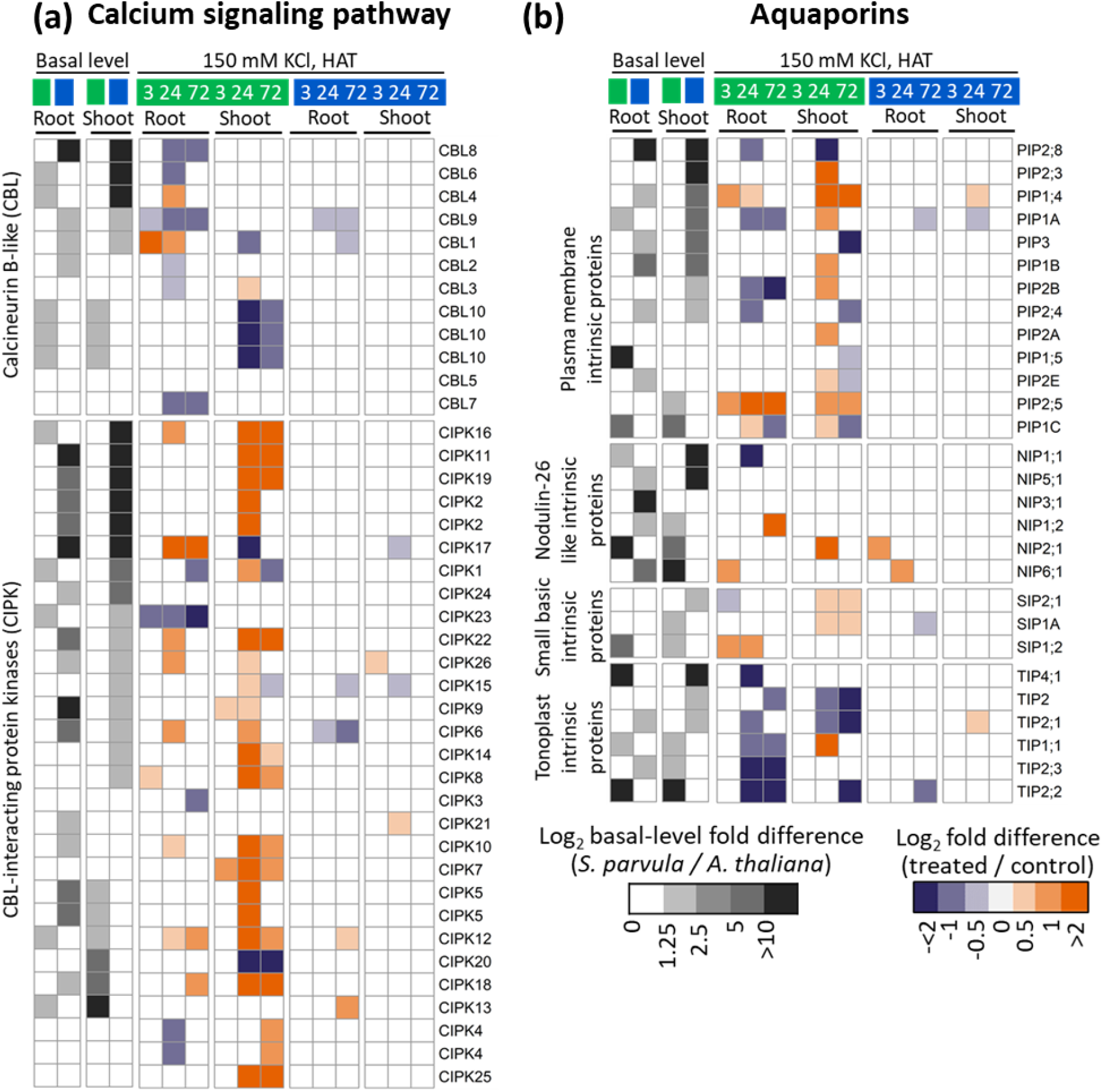
Genes coding for calcium signaling and aquaporins are differently regulated during high K^+^ stress. Genes associated with (a) calcium signaling pathway and (b) aquaporins during high K^+^ in root and shoot of *A. thaliana* and *S. parvula*. The significantly changed genes at least in one condition are presented in the heatmap. DEGs at each time point were called using DESeq2 compared to 0 h with a p-adj value based on Benjamini-Hochberg correction for multiple testing set to ≤0.01.

**Fig. S8.**
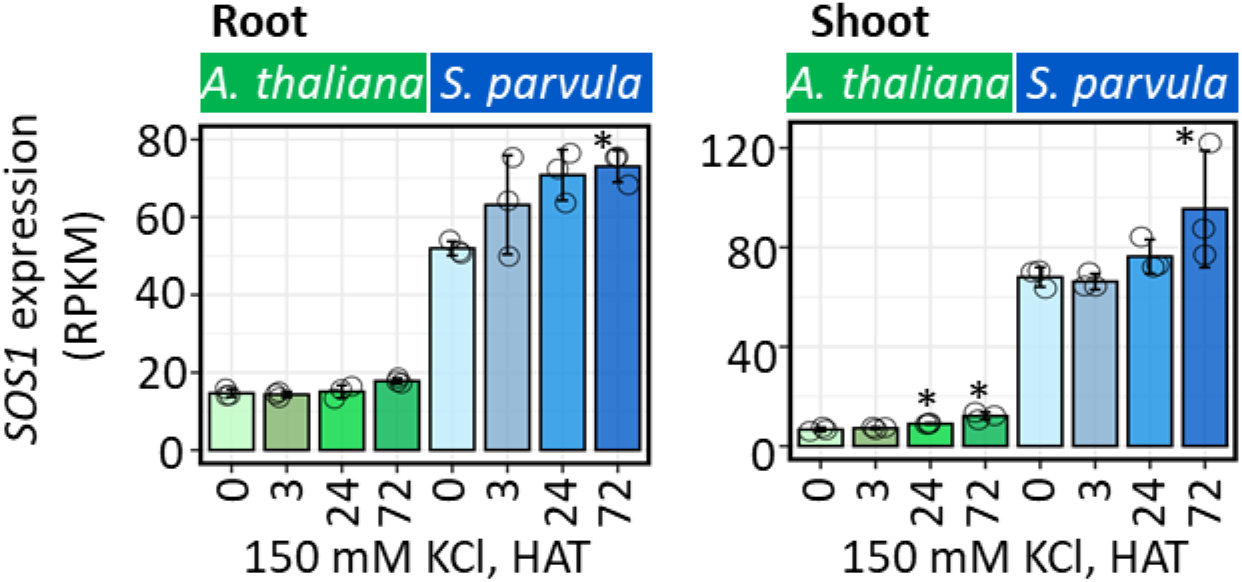
Normalized expression of *SOS1* in roots and shoots of *A. thaliana* and *S. parvula* under 150 mM KCl treatments. * represent DEGs at each time point called using DESeq2 compared to 0 h with a p-adj value based on Benjamini-Hochberg correction for multiple testing set to ≤0.01. Open circles indicate the measurement from each replicate.

## Supplementary table title

**Table S1** Relative abundance of metabolites for *A. thaliana* and *S. parvula* roots and shoots sample. 25-days-old hydroponically grown seedlings were treated for 24 and 72 hours after 150 mM KCl treatment and control samples were harvested together with the treated samples. Data are represented as the mean of at least 3 independent replicates. ≥5 plants per replicate were used.

**Table S2** Number of total reads and percentage of uniquely mapped reads to *A. thaliana* (TAIR10) or *S. parvula* v2.2 gene models for root and shoot transcriptomes under high K^+^. At-*A. thaliana,* Sp-*S. parvula*, C-control, 3-3 hours after treatment, 24-24 hours after treatment, 72-72 hours after treatment, R-root samples, S-shoot samples. 25-days-old hydroponically grown seedlings were treated for 3, 24, and 72 h after 150 mM KCl treatment. Control samples were harvested together with the treated samples. Data are represented as the mean of at least 3 independent replicates. ≥5 plants per replicate were used.

**Table S3** List of differentially expressed genes (DEGs) in 3, 24, and 72 hours after treatment (HAT) in *A. thaliana* and *S. parvula* root and shoot. DEGs were called using DESeq2 with a *p*-adj value based on Benjamini-Hochberg correction for multiple testing set to <0.01. Data are represented as mean of 3 independent replicates ± SD given using ≥ 5 hydroponically grown plants per replicate.

**Table S4** Enriched biological processes for a non-redundant set of induced and suppressed DEGs for *A. thaliana* roots and shoots sample under 150 mM KCl. DEGs were functionally annotated by a Gene Ontology (GO) enrichment test using BinGO in Cytoscape and enriched biological processes were further clustered based on shared genes using GOMCL with adj *p*-value <0.05 after false discovery rate correction with Benjamini-Hochberg correction.

**Table S5** Enriched biological processes for time point-specific DEGs (3, 3&24+24, 24&72, 72, and 3&24&72) for *A. thaliana* roots sample under 150 mM KCl. DEGs were functionally annotated by a Gene Ontology (GO) enrichment test using BinGO in Cytoscape and enriched biological processes were further clustered based on shared genes using GOMCL with adj *p*-value <0.05 after false discovery rate correction with Benjamini-Hochberg correction.

**Table S6** Enriched biological processes for time point-specific DEGs (3, 3&24+24, 24&72, 72, and 3&24&72) for *A. thaliana* shoots sample under 150 mM KCl. DEGs were functionally annotated by a Gene Ontology (GO) enrichment test using BinGO in Cytoscape and enriched biological processes were further clustered based on shared genes using GOMCL with adj *p*-value <0.05 after false discovery rate correction with Benjamini-Hochberg correction.

**Table S7** Enriched biological processes for time point-specific (3, 3&24, 24, 24&72, 72, and 3&24&72) induced and suppressed DEGs for *A. thaliana* roots sample under 150 mM KCl. DEGs were functionally annotated by a Gene Ontology (GO) enrichment test using BinGO in Cytoscape and enriched biological processes were further clustered based on shared genes using GOMCL with adj *p*-value <0.05 after false discovery rate correction with Benjamini-Hochberg correction.

**Table S8** Enriched biological processes for time point-specific (3, 3&24, 24, 24&72, 72, and 3&24&72) induced and suppressed DEGs for *A. thaliana* shoots sample under 150 mM KCl. DEGs were functionally annotated by a Gene Ontology (GO) enrichment test using BinGO in Cytoscape and enriched biological processes were further clustered based on shared genes using GOMCL with adj *p*-value <0.05 after false discovery rate correction with Benjamini-Hochberg correction.

**Table S9** Enriched biological processes for diametric responses (*i.e.* genes that are induced in one species when their orthologs are suppressed in the other species) in *A. thaliana* and *S. parvula* roots and shoots sample under 150 mM KCl. The pattern is extracted from Fig S6. DEGs were functionally annotated by a Gene Ontology (GO) enrichment test using BinGO in Cytoscape and enriched biological processes were further clustered based on shared genes using GOMCL with adj *p*-value <0.05 after false discovery rate correction with Benjamini-Hochberg correction.

**Table S10** Enriched biological processes for a non-redundant set of induced and suppressed DEGs for *S. parvula* roots and shoots sample under 150 mM KCl. The *S. parvula* DEG orthologs with *A. thaliana* were functionally annotated by a Gene Ontology (GO) enrichment test using BinGO in Cytoscape and enriched biological processes were further clustered based on shared genes using GOMCL with adj *p*-value <0.05 after false discovery rate correction with Benjamini-Hochberg correction.

**Table S11** Normalized gene expression clusters of ortholog pairs (OP) between *A. thaliana* and *S. parvula* in roots and shoots sample. Fuzzy K-means clustering was used to find temporally co-regulated clusters with a membership cutoff of >0.5. From initially identified 10 root and 11 shoot clusters, we filtered out clusters that did not show a response to K treatments in both species and identified 5 root (RC1, RC2, RC3, RC4, and RC5) and 3 shoot (SC1, SC2, and SC3) co-expression superclusters with distinct response trends. The orthologs from each cluster were functionally annotated by a Gene Ontology (GO) enrichment test using BinGO in Cytoscape and enriched biological processes were further clustered based on shared genes using GOMCL with adj *p*-value <0.05 after false discovery rate correction with Benjamini-Hochberg correction.

